# Vulnerability to memory decline in aging – a mega-analysis of structural brain change

**DOI:** 10.1101/2025.03.27.642988

**Authors:** Didac Vidal-Piñeiro, Øystein Sørensen, Marie Strømstrad, Inge K. Amlien, William Baaré, David Bartrés-Faz, Andreas M. Brandmaier, Gabriele Cattaneo, Sandra Düzel, Paolo Ghisletta, Richard N. Henson, Simone Kühn, Ulman Lindenberger, Athanasia M. Mowinckel, Lars Nyberg, Alvaro Pascual-Leone, James M. Roe, Javier Solana-Sánchez, Cristina Solé-Padullés, Leiv Otto Watne, Thomas Wolfers, the Vietnam Era Twin Study of Aging (VETSA), the Australian Imaging Biomarkers and Lifestyle flagship study of ageing (AIBL), the Alzheimer’s Disease Neuroimaging Initiative (ADNI), Kristine B Walhovd, Anders M. Fjell

## Abstract

Brain atrophy is a key factor behind episodic memory loss in aging, but the nature and ubiquity of this relationship remains poorly understood. This study leveraged 13 longitudinal datasets, including 3,737 cognitively healthy adults (10,343 MRI scans; 13,460 memory assessments), to determine whether brain change-memory change associations are more pronounced with age and genetic risk for Alzheimer’s Disease. Both factors are associated with accelerated brain decline, yet it remains unclear whether memory loss is exacerbated beyond what atrophy alone would predict. Additionally, we assessed whether memory decline aligns with a global pattern of atrophy or stems from distinct regional contributions. Our mega-analysis revealed a nonlinear relationship between memory decline and brain atrophy, primarily affecting individuals with above-average brain structural decline. The associations were stronger in the hippocampus but also spread across diverse cortical and subcortical regions. The associations strengthened with age, reaching moderate associations in participants in their eighties. While APOE ε4 carriers exhibited steeper brain and memory loss, genetic risk had no effect on the *change-change* associations. These findings support the presence of common biological macrostructural substrates underlying memory function in older age which are vulnerable to multiple age-related factors, even in the absence of overt pathological changes.

## Introduction

Episodic memory declines with age^1,2^, although individual trajectories vary significantly, with some experiencing marked cognitive decline and others maintaining relatively stable memory function over time^3^. Brain atrophy is considered a key contributor of these changes^4,5^. However, key questions remain poorly understood such as whether the *change – change* associations are dependent on age and genetic risk for Alzheimer’s disease (AD) or if the patterns are driven exclusively by a subset of individuals with severe atrophy. Further, extant research remains inconclusive on whether the effects reflect global patterns of brain atrophy^6,7^ or regional structural vulnerabilities, particularly in the hippocampus^8,9^. To examine these questions, we leveraged 13 datasets with more than 3,700 cognitively healthy adult participants who underwent repeated MRI scans and cognitive assessments together with simulations to guide results interpretation.

Both episodic memory and medial temporal lobe (MTL) structures tend to show relative stability across early adulthood and middle age in longitudinal studies, with a more marked decline from about 60 years^1,2,10–12^. In contrast, trajectories of cortical thickness outside the MTL tend to follow largely monotonic declines across adulthood, likely with subtle acceleration in thinning late in life^10,13–15^, and a relatively large degree of heterogeneity for the remaining subcortical structures^11,16,17^. Aging is also associated with increased interindividual variability in cognition and biological measures, including memory performance and brain structure^16,18^ and, even more importantly, the variability of brain and memory change also increases with age^19–21^.

Initial evidence for a relationship between memory function and brain structure in cognitively healthy aging came from cross-sectional research^22,23^, which indicated that differences in brain structure, particularly in the MTL, explained a modest amount of age-related variability in episodic memory function. Yet, it is now widely recognized that only longitudinal designs can effectively link co-occurring within-person changes in brain and cognition throughout the lifespan^24^. Indeed, longitudinal research has corroborated the association between memory decline and loss of grey matter in medial temporal features such as hippocampal atrophy^25–29^ and entorhinal thinning^8,30^. The findings align with the key role of these structures in episodic memory and their susceptibility to aging and AD^31–33^. Outside this region, associations have also been reported, albeit less consistently, in the frontal, parietal and temporal lobes^34,35^, as well as in global grey matter^6,27^. These results are consistent with the complex cortico-subcortical circuitry supporting episodic memory function^33,36–38^. Debate remains on whether these change-change associations are driven by a *main* factor of brain decline or by one or more of these memory-sensitive structures^6,26,29,34,39^. Current research shows evidence of both domain-general dimensions of cortical and cognitive change^6,40^ and domain-specific associations between MTL and episodic memory change^39^.

Since age affects both individual trajectories of brain structure and memory – and their variability across individuals - it is likely that age also moderates their relationship, such that the associations strengthen with increasing age. However, limited data exist on this topic and, when available, this data primarily concern the hippocampus. Cross-sectional evidence suggests stronger associations between hippocampus volume and memory performance in late life^41,42^. Using longitudinal data, Gorbach and colleagues^25^ found an association between hippocampal atrophy and memory decline that was significant in older (65 – 80 years) but not in middle-aged (55 – 60 years) individuals, and further suggested that steeper declines in memory and hippocampal volume facilitate detection of these associations.

The Apolipoprotein (APOE) ε4 allele represents the strongest known genetic risk factor for late-onset AD. In individuals with AD, ε4 carriers exhibit steeper brain atrophy, especially in limbic regions, and greater memory loss compared to non-carriers^43,44^. This pattern has also been observed in cognitively healthy older individuals^45,46^, although not universally^47^. Some theoretical models propose distinct AD subtypes based on the APOE ε4 allele^48^. In this account, the ε4 carrier subtype represents a more severe, limbic-dominant form of AD, characterized by steeper loss of memory and stronger links to amyloid pathophysiology. The non-carrier subtype is thought to represent a milder form, more heterogeneous, and more associated with environmental factors^49^. This model also predicts stronger associations between memory change and atrophy, particularly in the MTL, in ε4 carriers. Supporting this, some studies report a stronger brain-memory association amongst APOE ε4 carriers, in AD samples^50^ and, crucially, in a longitudinal sample of cognitively healthy older adults^9^.

Not all observed changes in brain structure in older individuals reflect long-term changes, i.e. *brain aging*, but also short-term variations due to known, e.g., physical or cognitive training programs^51,52^, and unknown factors, i.e., noise^20^. Hence, only a fraction of the observed changes occurring in brain structure over time may relate to memory decline. Thus, it is possible that most variation in brain structure over time is not *degenerative,* and that only individuals with severe atrophy show memory loss, leading to non-linear *change – change* associations.

Given the inherent heterogeneity in brain and cognitive trajectories, study samples, and analytical approaches – along with partially conflicting findings and the need to sample diverse populations for broad conclusions -, large-scale mega-analyses across cohorts are essential to accurately investigate change - change relationships between episodic memory and structural brain decline^5^. Here, we conduct such a study to address key questions in the field: 1) Do brain change-memory change associations become more pronounced with increasing age? 2) Are these relationships consistent with a global factor of brain structural decline, MTL vulnerability, or multiple regional contributions? 3) Are change-change associations more pronounced in at-risk individuals, namely those with above-average brain decline and/or carriers of the APOE ε4 allele? We used a normative modeling framework to harmonize 13 datasets with longitudinal MRI scans and cognitive assessments, resulting in a final sample of 3,737 cognitively healthy adults (10,343 MRI scans; 13,460 memory assessments) (**Table 1, Supplementary Figure 1**). We estimated individual change in memory (Δmemory) and 166 brain cortical and subcortical regions (Δbrain), with a particular focus on the hippocampus. We focused on thickness as a measure of cortical change given its susceptibility to age^53,54^, used a mega-analytical approach to maximize statistical power^55^, and general additive mixed models (GAMMs) to enable greater analytical flexibility. Still, for simplicity, throughout the text, we refer to individuals with above- and below-average brain decline – related to their age and sex peers – as *brain decliners* and *maintainers.* Finally, we provide complete statistics and visualizations in a **Supporting App** (https://vidalpineiro.shinyapps.io/brain_mem_change/) and simulate data to aid the interpretation of results.

**Table 1.** Main Sample sociodemographics. Main sociodemographic and observational detail of the main sample used in all main analyses. N = Total number of individuals or observations. NC = Non-carriers. C = Carriers. M = mean. SD = Standard Deviation. Obs. = Observations

| Dataset | Subjects | Apoe ε4 | Age |  | Obs. Memory |  |  | Obs. Brain |  |  | Time Brain |  | Time Memory |  |
| --- | --- | --- | --- | --- | --- | --- | --- | --- | --- | --- | --- | --- | --- | --- |
|  | N (m) | NC:C | M (SD) | range | N | M (SD) | range | N | M (SD) | range | M (SD) | range | M (SD) | range |
| adni | 223 (92) | 158:65 | 72.4 (6.3) | 55.8 – 89.9 | 1118 | 5.0 (2.7) | 2 - 13 | 984 | 4.4 (2.2) | 2 - 12 | 3.9 (2.1) | 1.6 – 9.6 | 4.6 (2.8) | 2.0 – 13.5 |
| aibl | 142 (70) | 94:78 | 71.5 (6.4) | 60.0 – 87.0 | 564 | 4.0 (1.1) | 2 - 5 | 493 | 3.5 (1.1) | 2 - 5 | 4.2 (1.6) | 2.0 – 8.0 | 4.9 (1.6) | 2.0 – 7.0 |
| bbhi | 256 (141) | 30:3 | 54.0 (7.2) | 41.2 – 66.1 | 512 | 2.0 (0) | 2 - 2 | 512 | 2.0 (0) | 2 - 2 | 2.4 (0.2) | 1.6 – 3.0 | 2.4 (0.2) | 1.6 – 3.0 |
| habs | 223 (94) | 157:63 | 73.3 (6.1) | 62.5 – 89.3 | 1180 | 5.3 (1.0) | 3 - 6 | 615 | 2.8 (0.6) | 2 - 4 | 4.4 (1.2) | 2.0 – 8.5 | 4.5 (1.1) | 2.0 – 8.5 |
| mpib | 214 (127) | 133:39 | 64.8 (15.0) | 24.5 – 83.1 | 572 | 2.7 (0.5) | 2 - 3 | 428 | 2.0 (0) | 2 - 2 | 2.3 (0.4) | 1.5 – 3.1 | 4.5 (1.5) | 1.6 – 6.4 |
| oasis3 | 431 (171) | 287:137 | 67.2 (8.9) | 43.5 – 95.6 | 3502 | 8.1 (4.0) | 2 - 23 | 1458 | 3.4 (1.5) | 2 - 8 | 6.4 (3.4) | 1.5 – 15.8 | 9.9 (4.2) | 2.0 – 24.0 |
| ous | 93 (42) | 54:38 | 73.2 (6.0) | 64.7 – 90.0 | 596 | 6.4 (1.0) | 3 - 7 | 366 | 3.9 (1.3) | 2 - 6 | 6.0 (2.6) | 1.8 – 9.5 | 5.8 (0.9) | 2.1 – 6.9 |
| preventad | 184 (53) | 115:69 | 63.6 (5.0) | 55.1 – 83.3 | 874 | 4.8 (1.0) | 3 - 6 | 1061 | 5.8 (1.0) | 4 - 7 | 3.3 (0.8) | 1.9 – 4.7 | 3.2 (0.8) | 1.6 – 4.5 |
| ub | 77 (28) | 67:10 | 68.6 (5.0) | 51.7 – 78.1 | 214 | 2.8 (0.4) | 2 - 3 | 208 | 2.7 (0.5) | 2 - 3 | 3.5 (1.0) | 1.6 – 5.0 | 3.7 (1.0) | 1.6 – 5.2 |
| uio | 304 (120) | 100:72 | 47.9 (19.1) | 20.0 – 85.5 | 781 | 2.6 (0.7) | 2 - 4 | 799 | 2.6 (0.8) | 2 - 7 | 6.2 (2.8) | 2.4 – 11.5 | 6.0 (2.7) | 2.4 – 11.5 |
| ukb | 106 6 (525) | 775:272 | 62.3 (7.0) | 47.0 – 79.5 | 2132 | 2.0 (0) | 2 - 2 | 2132 | 2.0 (0) | 2 - 2 | 2.3 (0.1) | 2.0 – 2.7 | 2.3 (0.1) | 2.0 – 2.7 |
| umu | 47 (27) | 26:15 | 43.8 (12.8) | 25.0 – 75.0 | 94 | 2.0 (0) | 2 - 2 | 94 | 2.0 (0) | 2 - 2 | 4.3 (0.4) | 4.0 – 5.0 | 4.3 (0.4) | 4.0 – 5.0 |
| vetsa | 477 (477) | 354:122 | 58.0 (3.9) | 51.1 – 70.1 | 1321 | 2.8 (0.4) | 2 - 3 | 1193 | 2.5 (0.5) | 2 - 3 | 9.3 (2.7) | 4.5 – 13.4 | 10.0 (2.4) | 4.5 – 14.4 |
| all | **3737 (1967)** | **2350:953** | **62.5 (11.6)** | **20.1 – 95.6** | **13460** | **3.6 (2.6)** | **2 - 23** | **10343** | **2.8 (1.3)** | **2 - 12** | **4.5 (3.0)** | **1.5 – 15.8** | **5.1 (3.6)** | **1.6 – 24.0** |

## Results

### Association between brain change and memory change. Main effects

We used GAMMs to assess the relationship between brain change (Δbrain) (cortical thickness and subcortical volume) and memory change (Δmemory). Δbrain was modelled as a smooth term and dataset as random intercept. Sex and age trends were regressed out during normative modelling-based preprocessing and thus not included in the higher-level model. Henceforth, change in brain and memory for a given individual is relative to their age and sex-peers. Control analyses with age, sex, or intracranial volume (ICV) as covariates, or with random slopes per dataset did not substantially affect the outcome of the main results (**Supporting App**). Data were weighted to account for differences in reliability of change, as longitudinal data with fewer observations and shorter follow-up time contain more uncertainty^20^. Regions were defined based on the *Destrieux*^56^ cortical and *aseg*^57^ subcortical atlas within FreeSurfer. Nineteen regions showed significant, False Dscovery Rate (FDR) corrected (p_FDR_ < 0.05) (**Figure 1a**), mostly subcortical structures and temporal regions. The relationship for all these regions was non-linear, generally showing an association between Δbrain and Δmemory only when Δbrain was steeper than average (i.e. in *brain decliners*). When brain decline was milder than average (i.e. in *brain maintainers*), the association between Δbrain and Δmemory disappeared. Left and right hippocampus (β_weighted[w]_[l] = .168, pFDR < 0.001); β_w_[r] = .168, p_FDR_ < 0.001), left amygdala (β_w_ = .155, p_FDR_ < 0.001), left thalamus (β_w_ = .135, p_FDR_ = 0.03), right long insular gyrus (β_w_ = .135, p_FDR_ = 0.02) and left parahippocampal gyrus (β_w_ = .131, p_FDR_ = 0.02) were amongst the regions showing strongest change – change associations in above-average brain decliners. See **Figure 1b** for visualization of selected regions. See **Supplementary Table 1** for statistics in significant regions. See **Supporting App** for complete statistics and visualization in all regions. No strong evidence for left – right asymmetry in change – change associations was found (**Supplementary Figure 2; SI**).

**Figure 1.**
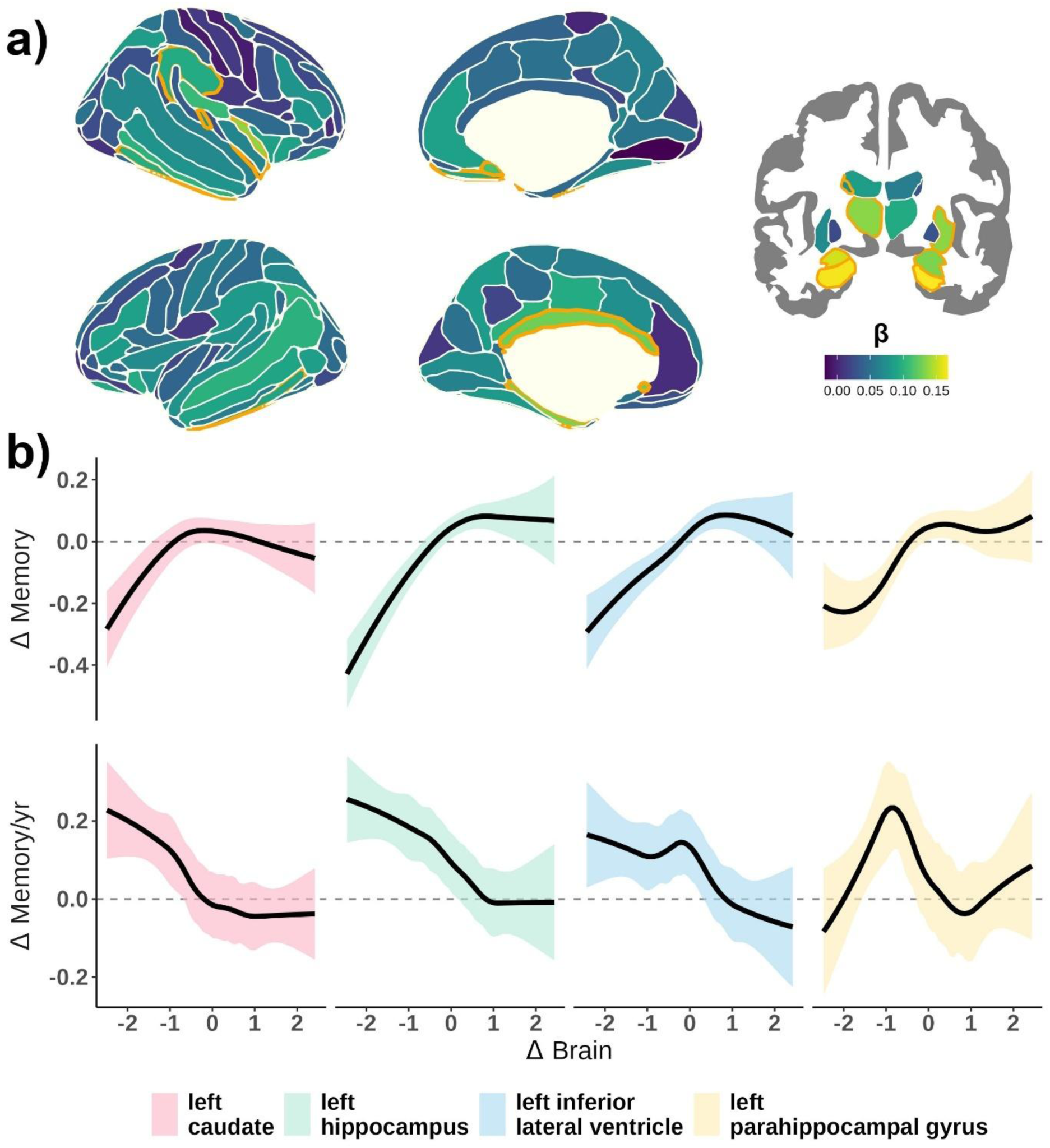
Memory change – brain change associations. **a)** Estimates (β_w_) for associations between Δbrain and Δmemory with orange color line representing p_FDR_ < 0.05. Δbrain represents atrophy in subcortical and thinning in cortical regions. Weighted betas were estimated as the density-weighted mean correlation between Δbrain and Δmemory, using the derivative function. **b)** Change – change association for selected regions. Upper plots display the smooth associations while lower plots show their derivative (i.e. association between Δbrain and Δmemory for each Δbrain value). Shaded ribbons represent 95% CIs. See **Supplementary Table 1** and **Supporting app** for more information.

### Association between brain change and memory change. Age interactions

Next, we assessed whether the association between Δbrain and Δmemory varied with increasing age using tensor smooths (i.e. interactions between marginal smooth terms) as implemented in GAMM. For 7 regions, age significantly moderated the change - change associations (p_FDR_ < .05) (**Figure 2a**) namely left (p_FDR_ = 0.02) and right hippocampus (p_FDR_ = 0.02), right inferior lateral ventricle (p_FDR_ < 0.001), left lateral ventricle (p_FDR_ = 0.05), right caudate (p_FDR_ = 0.03), right putamen (p_FDR_ = 0.03), and left short insular gyrus (p_FDR_ = 0.02). In most of these regions we found that change – change associations increased with higher age and progressively included *brain maintainers*. Change – change associations in some regions begin to be apparent between 50 and 60 years. These regions differ in the specific *shape* of the interaction. For example, change – change associations in brain decliners are first apparent at ≈50 years in the right hippocampus, at ≈60 years for the left hippocampus and the right lateral inferior ventricle and at ≈70 years for the short insular gyrus and the right caudate. Similarly, associations between Δbrain and Δmemory in brain maintainers are apparent from ≈70 years in the left hippocampus and the right lateral inferior ventricle but not in other regions such as the short insular gyrus or the right hippocampus. We use the left hippocampus to illustrate these effects: the relationship between Δbrain and Δmemory in brain decliners is β_w_ = -.04 at age 40 years, β_w_ = .02 at 50 years, β_w_ = .13 at 60 years, β_w_ = .23 at 70 years, and β_w_ = .29 at 80 years. In contrast, the relationship between Δbrain and – Δmemory in brain maintainers is non-existent until age 70 (β_w_ = .13), with a small increase at age 80 (β_w_ = .19). See **Figure 2b** for visualization of selected regions. See **Supplementary Table 2** for statistics in significant regions and complete statistical outcomes in the **Supporting App**.

**Figure 2.**
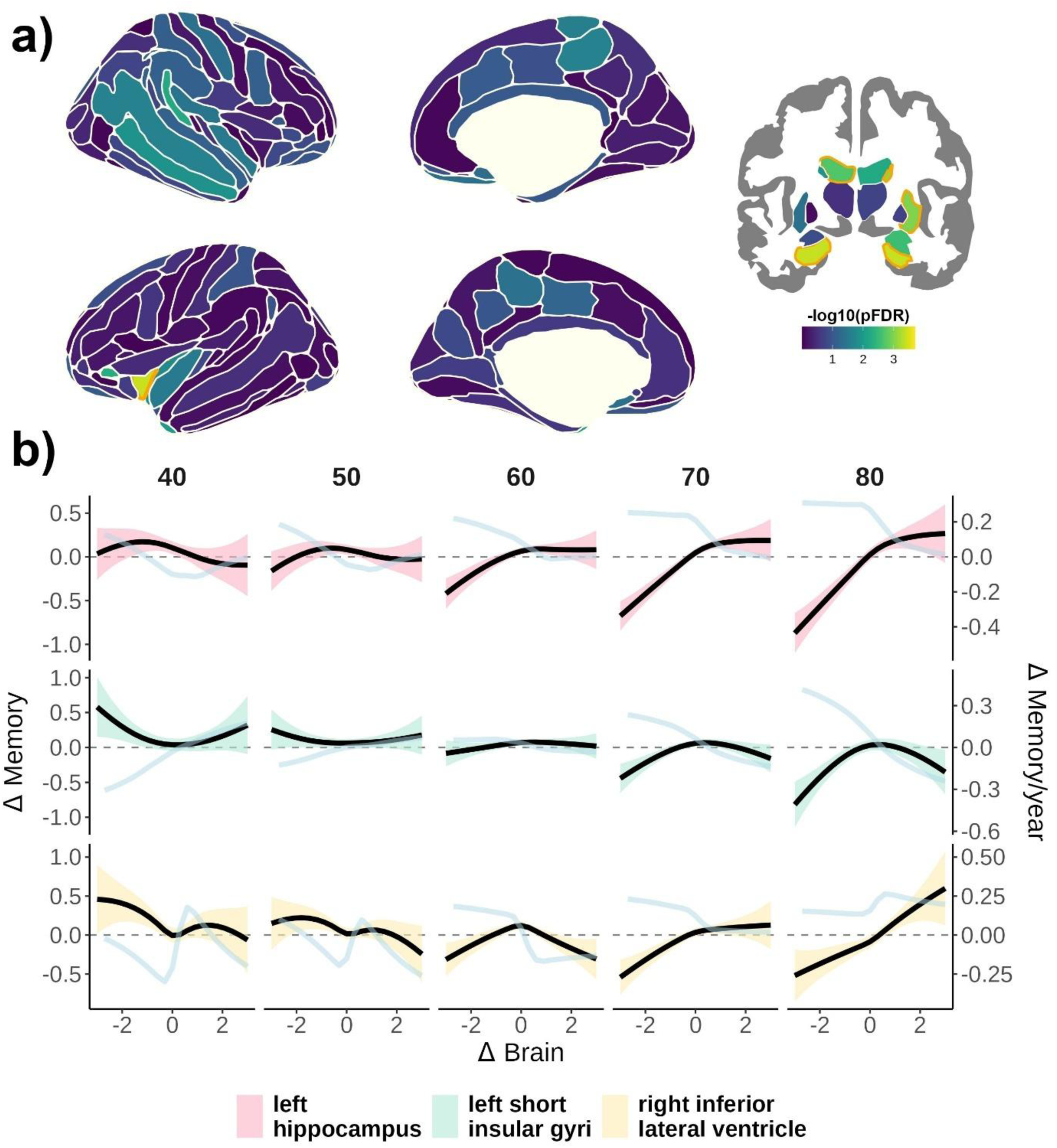
Effect of age on Memory change – brain change associations. **a)** −log10(p_FDR_) values of the effect of age on Δbrain – Δmemory associations modeled as GAMM-based tensor interaction terms. Orange color line represents p_FDR_ < 0.05. Δbrain represents atrophy in subcortical and thinning in cortical regions. **b)** Change – change association for selected regions at specific ages. Black line - and colored 95% CIs ribbons - display Δbrain - Δmemory associations at a given age. The lightblue line represents their derivative (i.e. association between Δbrain and Δmemory for each Δbrain value). See **Supplementary Table 2** and **Supporting app** for more information. Note that both Δbrain is derived from normative data and thus does not necessarily reflect the same amount of decline at each age.

### Dimensionality of brain change

Next, we explored the dimensionality of those regional brain changes associated with memory decline. Is memory loss associated with a single global effect of brain decline, or does it reflect region-specific contributions? We computed the correlation of brain change across brain regions (**Figure 3a**, mean r = .14 [0.10]; range = -.04 - .58) and carried out a PCA and a consensus clustering analysis to investigate this question. On the one hand, the PCA revealed that the first principal component (PC1) accounted for – a somewhat modest - 20.7% of the total variance, with all its loadings pointing in the same direction, and a significant, ≈two-fold, fall in the variance explained by subsequent components (**Figure 3b**). This suggests a pattern of brain change that to some degree aligns with the presence of a global pattern of brain decline.

**Figure 3.**
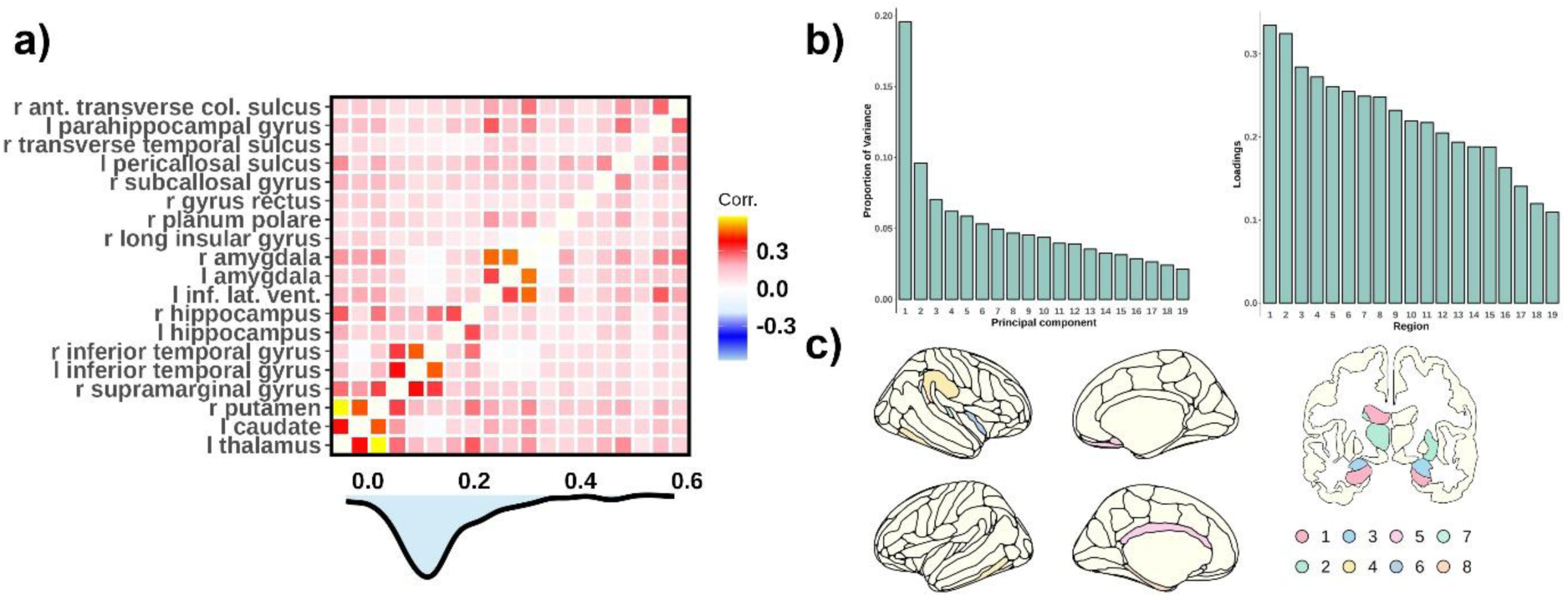
Dimensionality of brain change. a) Cross-correlation matrix of brain change across regions with significant (p_FDR_ < .05) Δmemory – Δbrain associations. Below, a density plot of the correlation coefficients. b) Variance explained by the principal PCA components and loading of the first component on the brain regions showing significant Δbrain – Δmemory associations. See ID to region name correspondence in **Supplementary Table 3**. c) Optimal consensus clustering solution (K = 8 clusters).

On the other hand, consensus clustering analysis was performed to explore whether the effects were regional. Importantly, we tested using Monte-Carlo simulations whether the clustering solution rejected the null hypothesis of K = 1 cluster. Several clustering solutions rejected the null hypothesis, with 8 clusters being the best solution (**Figure 3c; Supplementary Figure 3, Supplementary Table 3**). Three clusters were subcortical (clusters #1 - #3) - one comprised of the left and right hippocampus and the left lateral ventricle - and 5 were cortical. Further, we assessed whether change in any of these clusters was associated with memory change, by respectively controlling for the hippocampus-based cluster, the main factor of brain decline, and by introducing all clusters together in a single model. Five clusters showed significant Δbrain – Δmemory associations controlling for the hippocampal-based cluster (cluster #1): cluster #3, the left and right amygdala (β_w_ = .11, p = 0.01); cluster #5, left pericallosal sulcus, right gyrus rectus, and right subcallosal gyrus (β_w_ = .16, p = 0.002), cluster #6, right long insular gyrus and planum polare (β_w_ = .06,p = 0.01); cluster #7, right temporal transverse sulcus (β_w_ = .05,p = 0.02); and cluster #8, left parahippocampal gyrus (β_w_ = .05, p = .03). Similar results were found when using the main component of brain decline and when all clusters were added in a single model. See **SI** and **Supplementary Table 4** for detailed information. Altogether, the results suggest both global and regional factors influence the associations between brain and memory change.

### Association between brain change and memory change. APOE ε4 effects

A total of 3,149 subjects had APOE data available. Of these, 27.8 % were carriers of the APOE ε4 allele (carriers vs. non-carriers). First, we assessed whether carriers of the APOE ε4 allele showed steeper decline in brain or memory and whether this relationship was associated with age. For the main effects, we used linear mixed models with APOE ε4 allele as predictor and dataset as random intercept. For the interaction, we used GAMM with age as a smooth term by APOE ε4 allele as an ordered factor. APOE ε4 was not significantly related to memory decline (β = -.035, t[p] = −0.95[0.34]) (**Figure 4a**) but the relationship between APOE status and memory change increased with age (p = 0.03), that is, APOE ε4 carriers showed less memory decline until ≈60 years of age, and more memory decline thereafter (**Figure 4b**).

**Figure 4.**
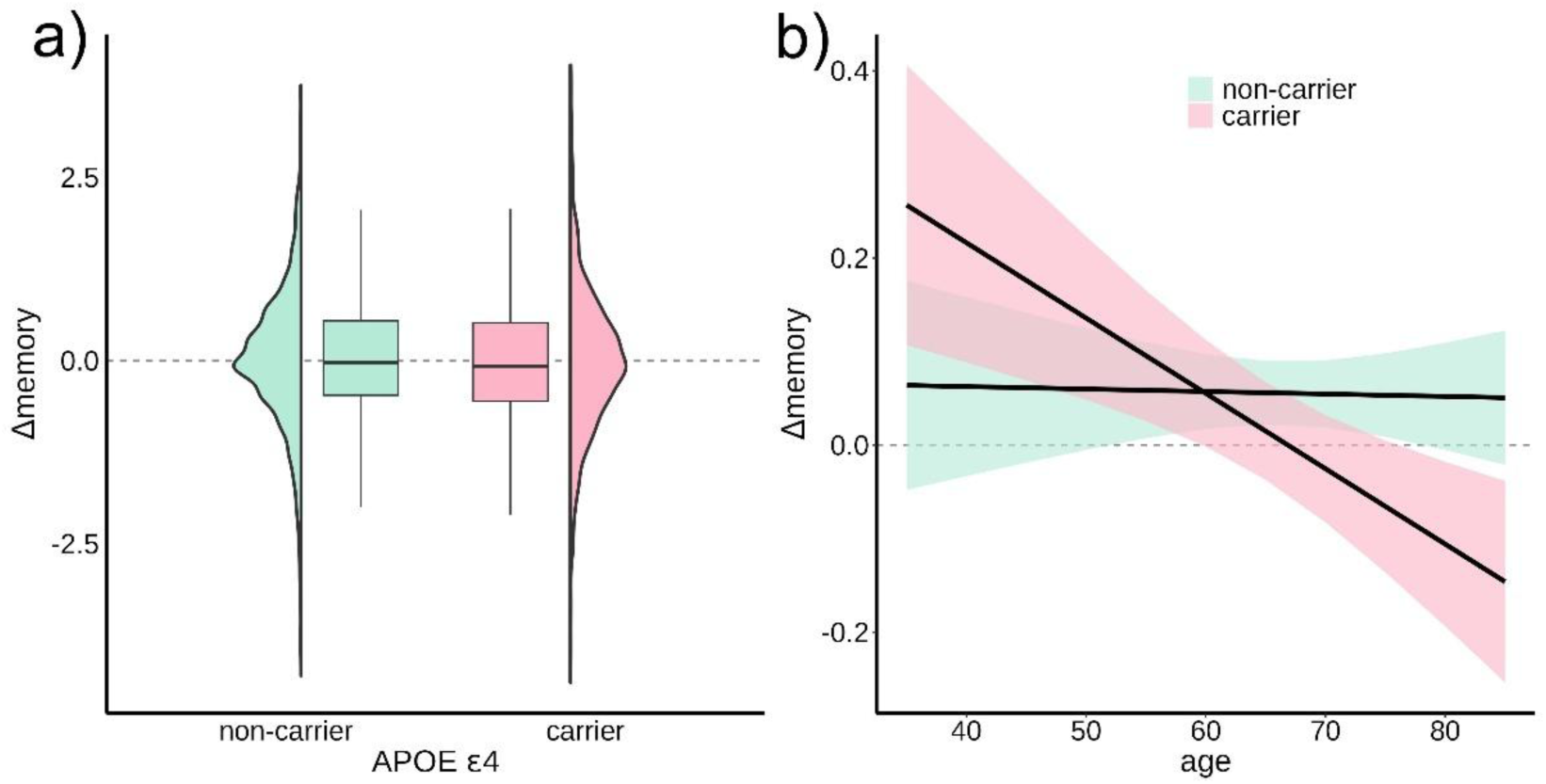
APOE ε4 associations with memory change. a) Association between APOE ε4 (carriers, non-carriers) and memory change. b) Association between APOE ε4 (carriers, non-carriers) and memory change as a function of age. Ribbons represent 95% CI.

APOE ε4 was associated with steeper left and right hippocampal atrophy (β_l_ = -.135, p_FDR_ = 0.004; β_r_ = -.138, p_FDR_ = 0.003) as well as right amygdala atrophy (β = -.154, p_FDR_ < .001) (**Figure 5a**). Considering all regions together, no global effect of the APOE ε4 allele on brain decline was found (β = -.004, t = −1.30, p = .20) (**Figure 5b**). APOE ε4 was not significantly associated with steeper brain decline (p_FDR_ > .10) with higher age. However, at an uncorrected level, APOE ε4 associated with a higher degree of hippocampal atrophy with increasing age (p = 0.01, 0.04, respectively) (**Supplementary Figure 4**).

**Figure 5.**
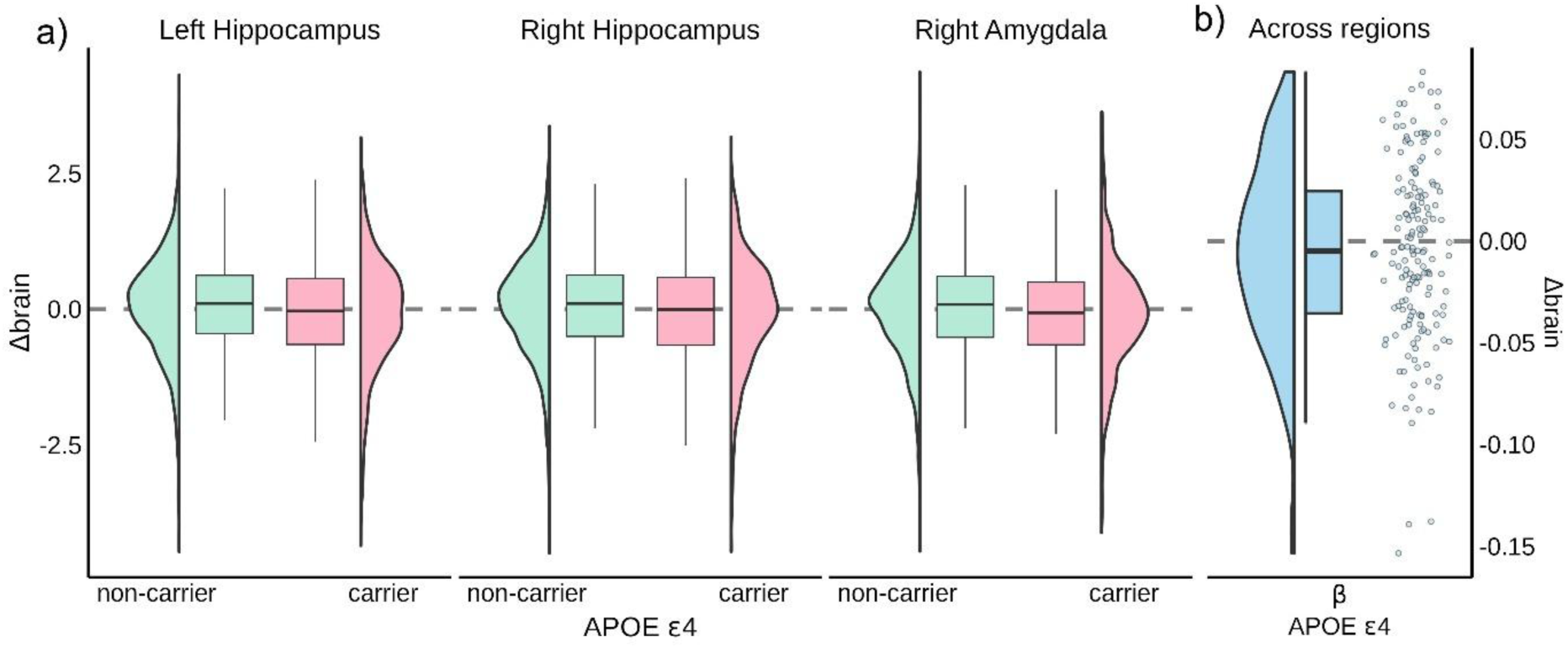
APOE ε4 associations with brain change. a) Association between APOE ε4 (carriers, non-carriers) and brain change in the left and right hippocampus, and the right amygdala (p_FDR_ < 0.05). b) Effect of APOE ε4 on brain change across all regions. Each point represents a region. Note that three regions with more negative effects of APOE ε4 correspond to those displayed in panel a).

Next, we assessed whether being an APOE ε4 carrier had any influence on the association between Δmemory and Δbrain. No region showed moderating effects of APOE ε4 on Δbrain – Δ memory associations (p_FDR_ > .50). APOE ε4 did not significantly moderate the relationship between Δbrain and Δmemory in the left and right hippocampus (p[l]_unc_ = .44, p[r]_unc_ = .61) (**Figure 6a**). Finally, no regions showed a significant interaction between APOE ε4, age, and Δbrain on Δmemory (p_FDR_ > .15). Left and right hippocampus showed comparable Δbrain - Δmemory associations with age regardless of APOE ε4 status (p[l]_unc_ = .66, p[r]_unc_ = .66) (**Figure 6b**). When using a sample of only APOE ε4 non-carriers, the *change – change* effects and the moderator effects of age were comparable to those for the main sample (**SI**).

**Figure 6.**
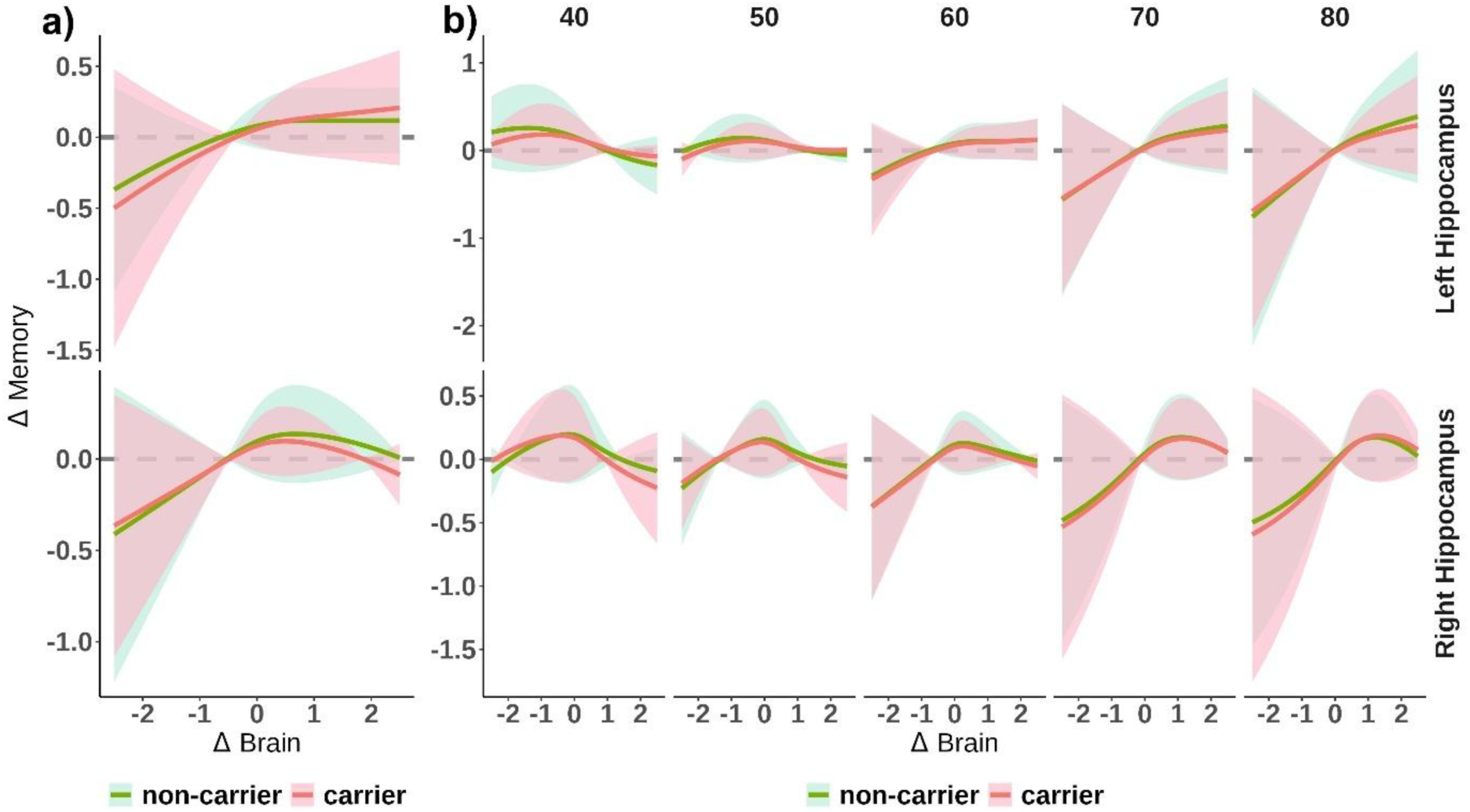
APOE ε4 effect of brain change – memory change associations. a) Δbrain - Δmemory associations as a function of APOE ε4 (carriers vs. non-carriers) for the left and right hippocampus. b) Δbrain - Δmemory associations as a function of APOE ε4 (carriers vs. non-carriers) and age for the left and right hippocampus. None of the terms were significant (p > 0.40).

See complete statistics and additional visualization for all APOE ε4 analyses in the **Supporting App**. Altogether, the results show that cognitively healthy carriers of the APOE ε4 allele have steeper rates of brain and memory decline, specifically in old adulthood, but no evidence of stronger Δbrain – Δmemory associations. The regional associations between brain change and memory change exist independently of an increased presence of pathological processes, and of cognitive changes associated with genetic risk of AD.

### Mechanisms behind brain change and memory change associations. A post-hoc simulation analysis

Finally, we aimed to provide potential explanations for the empirical findings, focusing on the non-linear *change – change* associations, the moderating effect of age, and the absence of APOE ε4 effects. We used a simplified schematic model using the *sn* package^58^ where *observed* brain data was the result of two underlying sources. The first source represented *brain aging,* characterized by a negatively skewed distribution with mean decline indicative of long-term, *degenerative* changes. This component is universal, as most individuals exhibited some degree of decline, with negative skewness arising from a subset of individuals undergoing accelerated brain aging^16,20,22,32,59^. The second source was modeled as a Gaussian distribution centered around zero, representing measurement error and other short-term influences^20,51,52^. Memory decline was modeled as linearly related to the *brain aging* component, plus random Gaussian noise. To explore the moderating effects (or lack thereof) of age and APOE ε4, we adjusted the parameters of the *brain aging* component, including mean, dispersion (e.g. variability), and skewness.

The simulation results revealed that *observed* brain decline was non-linearly associated with memory decline, with the relationship flattening among *brain maintainers* (**Figure 7a**) mimicking the empirical *change – change* associations. Increasing the variance of the *brain aging* component strengthened the *change – change* associations and affected *brain maintainers*, replicating the moderating effect of age (**Figure 7c**). Conversely, increasing mean decline and skewness did not alter the *change – change* associations despite leading to steeper mean decline in both brain and memory measures (**Figure 7b, d**). See details in **SI**.

**Figure 7.**
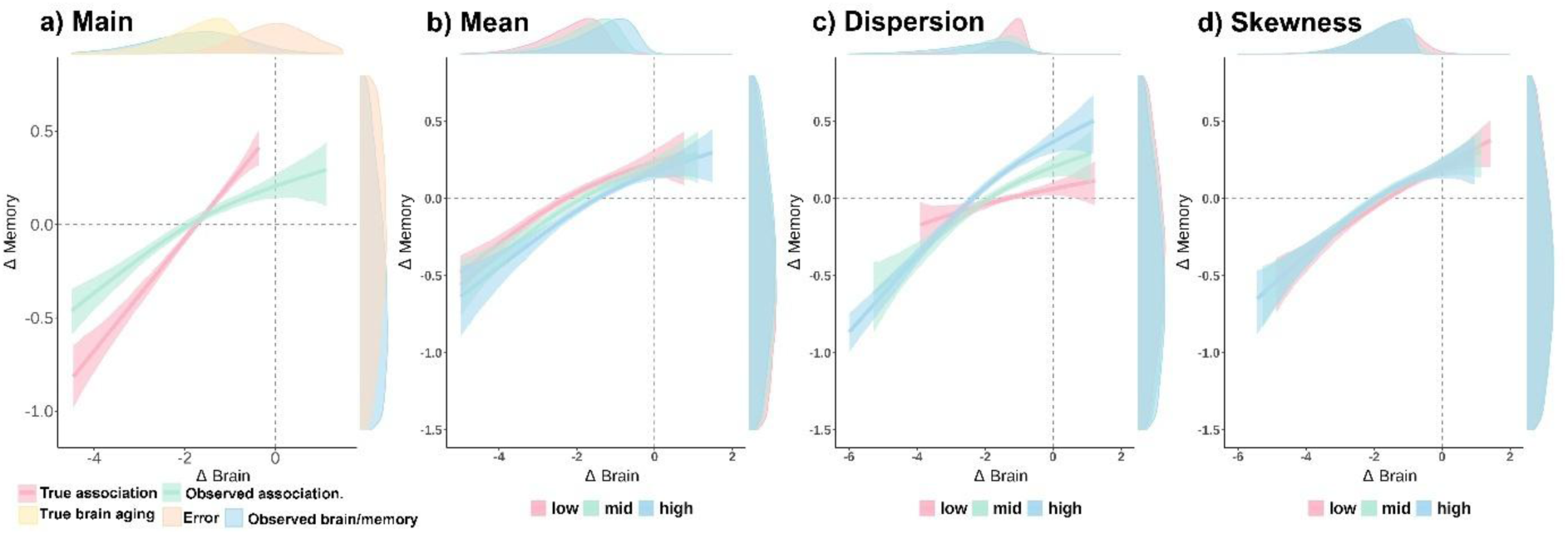
Theoretical basis for brain change – memory change associations. a) The main simulations demonstrate how a skewed true (latent) distribution of brain decline, which has a linear association with memory decline (red line) results in non-linear observed associations between observed (measured) brain decline and memory decline (green line) due to the measurement noise. Density plots illustrate different distributions: yellow represents true brain and memory decline, orange measurement error, and blue observed brain and memory. b-d) The impact of distribution moments on true brain distribution on observed Δbrain – Δmemory associations. Specifically, the effect of b) mean decline, c) dispersion, and d) negative skewness. Density plots correspond to the underlying distribution of true brain and memory decline. Ribbons indicate the 95% CIs across N = 1000 simulations. GAMs were used to fit the associations. See **SI** for more details.

## Discussion

By mega-analyzing data from over 3700 cognitively healthy adults and 13 independent longitudinal studies, we found that changes in brain structure are associated with changes in episodic memory across several cortical and subcortical areas, with the strongest associations in the MTL. These associations became more pronounced with increasing age, while no evidence was found for stronger relationships in APOE ε4 carriers. We argue for common macrostructural systems supporting memory function, where multiple factors converge to increase vulnerability in older age. The implications of these findings are discussed below.

### Brain decline – Memory loss associations; a generalized phenomenon? The case of above-average decliners

Observed brain decline is non-linearly associated with memory decline, with stronger associations in *brain decliners*, i.e. individuals exhibiting above-average brain decline relative to their age and sex, compared to *brain maintainers.* This novel finding within the context of cognitively healthy aging differs from previous research, which may have over relied on linear regression models and modest sample sizes. At first sight, it suggests *change – change* associations are constrained to a specific population of individuals with steeper brain decline, at-risk for pathological neurodegeneration. Yet, our simulations challenge this interpretation rather suggesting the non-linear trends result from the presence of multiple sources contributing to noisy measures of brain change. Amongst these, one component, i.e. brain aging, has linear associations with memory decline and is characterized by a negative mean and skewness. These assumptions align well with current evidence, including, critically, the skewed distribution of brain aging ^16,20,22,32,59,60^. The findings suggest that changes in *brain aging* are dimensional, skewed, and inherent process, which is an important determinant of memory loss in cognitively unimpaired elderly. Rather than a categorical view, where a *degenerative* component of brain change is limited to vulnerable individuals, it is the combination of noise and a skewed distribution that limits our ability to observe empirical associations in individuals with less brain change. These results likely underpin other categorical distinctions in neurocognitive aging^12^.

### Brain decline – Memory loss associations; a generalized phenomenon? The case of APOE ε4 carriers

APOE ε4 is associated with steeper brain decline and memory decline but does not affect the *change – change* associations between brain structure and memory. Carrying the APOE ε4 allele is the strongest genetic risk for sporadic AD, with dose-dependent effects^61,62^. Older carriers of the APOE ε4 allele exhibited steeper memory and brain decline, particularly in the hippocampi and the right amygdala, aligning with many other studies^45,46^. It is plausible that a higher proportion of APOE ε4 carriers are on a path to clinical disease manifestation of AD, putatively driven by the spreading of Tau deposition – which is strongly linked to steeper brain atrophy, memory decline, and short-term clinical diagnosis^63^. However, the *change – change* associations and the moderating effects of age were not influenced by APOE ε4 status, nor are they likely affected by *preclinical AD*. First, in some regions, the *change – change* associations were evident before age 60, when prevalence of Tau deposition is generally very low^63,64^. Aβ deposition before age 60 is slightly more common, but when controlling for Tau, the influence of Aβ on brain and memory decline is modest at best^63,65–67^. Second, the associations are not constrained to the MTL, where earlier preclinical changes are observed in AD. Third, if preclinical AD were to influence the Δbrain - Δmemory associations, APOE ε4 non-carriers would display attenuated or age-delayed associations between brain and memory decline. One study using linear models reported stronger associations between hippocampal change and memory decline in cognitively healthy APOE ε4 carriers^9^, arguing that APOE ε4 carriers had a more hippocampal-centric pattern of atrophy in line with categorical aging and disease models. The current results align with a more hippocampal-centric pattern of atrophy, but also fit well with a dimensional view of aging, where APOE ε4 contributes to accelerated brain aging, without changing the mechanisms underlying the *change – change* associations. This is captured by the simulation analyses, which illustrates that a *brain aging* distribution with either steeper mean decline or higher skewness in APOE ε4 carriers would lead to both steeper brain and memory decline but similar strength of the *change – change* associations. Altogether, these results fit with a dimensional view of aging, where APOE ε4 and early preclinical change in AD are one of many pathways affecting *common* biological substrates that determine memory function in older age, namely regional and global macrostructural atrophy.

### Brain decline – Memory loss associations; a generalized phenomenon? The case of age

Age strengthens the associations between brain decline and memory decline, gradually extending to *brain maintainers* across most significant regions. The hippocampi are among the earliest regions to exhibit these associations, emerging in the late fifties. This finding aligns with previous indirect evidence that the relationship between brain and memory decline strengthens with age^25,41,42^. Simulated data identified dispersion - greater variability in brain change across individuals - as the key factor driving stronger associations with age. This aligns with prior research indicating increased variability in both brain levels and brain change^16,20^, a pattern also observed in cognition, including episodic memory^18,19^. Interestingly, mean decline did not significantly affect the strength of the brain - cognition associations. The steeper memory and MTL declines from around 60 years of age^10,11,15,16^ are therefore not direct causes of these associations but rather serve as indirect markers. Since *brain aging* follows a unidirectional trajectory - where everyone experiences some degree of decline over time - greater variability in *brain aging* gives rise to steeper rates of brain decline. Overall, age is the primary determinant of *degenerative* brain change, and henceforth of change – change associations. Below a certain age, *brain aging* – or better said, population-level variability – is minimal, making it unlikely to be a key factor behind episodic memory loss; if a meaningful decline in episodic memory occurs in young adulthood^1,2,68^. What makes age the prime risk factor for brain decline and which age-related factors may explain variations in brain (and memory) decline, remain amongst the most critical questions in the field. All points out towards a multidimensional view, were brain systems, even in the absence of overt pathological changes, are highly vulnerable to several aging factors^32^.

### Regional associations between brain change and memory change

Hippocampal atrophy unsurprisingly showed the strongest associations with memory decline over time. This is consistent with earlier studies in the context of cognitively healthy elderly^25–29^, the key role of hippocampus in episodic memory^33,37^, and its vulnerability to aging^32^. In contrast, ventricular associations likely reflect global, non-specific patterns of brain atrophy and have also been reported elsewhere^27,34^. The mechanisms underlying observed caudate, thalamus, putamen, and amygdala *change – change* associations remain unclear and require experimental approaches to move beyond speculation. One possibility is altered coupling between these regions and the hippocampus, as they all exhibit connectivity changes with the hippocampi during aging and episodic memory tasks^69–72^. Associations between cortical thinning and memory decline were weaker than those observed for subcortical structures^29^. Cortical thinning was chosen due to its high sensitivity to change; however, this sensitivity may render it more susceptible to influences unrelated to aging and long-term memory decline. Among the regions surviving multiple comparisons correction, the left parahippocampal gyrus stood out. It’s anterior portion encompasses the entorhinal cortex, which serves as the main interface between the neocortex and the hippocampus, and is critically involved in memory^37,73,74^. Six additional regions in the temporal lobe were associated with memory decline, likely reflecting their roles in auditory, visual, or multimodal processing and integration. Most of the remaining regions, such as the pericallosal sulcus, the supramarginal gyrus, and the long insular gyrus, pertain to an action-mode network involved in task-positive, goal-directed behavior^75^. The overall pattern consists of relatively higher-order regions associated with both goal-directed and internal self-referential processing which manifest the particular requirements that memory encoding and retrieval impose on the brain, that is orchestrating dynamics amongst – often antagonistic - large-scale networks^76^.

### Dimensionality of brain change: Global decline or regional contributions?

Previous research has suggested the associations between brain atrophy and memory decline are driven either by a global factor of brain decline or are constrained to the MTL^6,25,39^. Our findings partially support both views, as we found evidence for a global factor of brain decline, while hippocampal atrophy showed the strongest regional associations with memory decline. However, the results reveal a more nuanced picture, with evidence for clustering observed. While the clustering solution made both topological and functional sense, caution is warranted regarding the specific solution obtained, as multiple solutions were plausible, and the input data were selected based on the somewhat arbitrary criterium of statistical significance. Nonetheless several clustering solutions outperformed the one-factor solution with some of these remained associated with memory decline even after controlling for hippocampus or global decline. Most existing research, including this study, does not fully disentangle cross-regional correlations in *brain aging* from correlated errors. In any case, the current results indicate that a decline in regions critical to lower-order functions, such as attention, or indirectly related to memory via reward or executive control systems, contribute to memory loss independently of the integrity of medial lobe structures. Note also that the relationship between some observed regions and memory decline may be explained by change in global cognition^39^.

### Technical considerations and limitations

The study required considerable analytical flexibility, which may influence the outcome. A multiverse approach^77^ was unfeasible due to data availability, technical, and computing constraints. Key considerations include: I) <u>Normative modeling-based normalization</u> using Hierarchical Bayesian Regression^78,79^, a flexible technique that often outperforms other harmonization methods^80^. It standardizes Z-scores based on age and sex, making data relative and somewhat challenging to interpret. However, it eliminates most age-related homoscedasticity and facilitates comparisons with other research, since data are aligned to an openly available norm. II) <u>Bootstrapped p-values</u> were estimated to better control the false positive rate in GAMMs^81^, leading to somewhat reduced power compared to linear models when the observed relationship is linear, although it can be argued that it hardly ever is. <u>III) Estimation of change scores</u> via linear changes over time per individual, with <u>weighting</u> applied to control for differences in longitudinal reliability as individuals with longer follow-up times and more observations contribute with more reliable data^20,82^. This represents a compromise choice, balancing data quality, flexibility, and interpretability; yet other approaches based, e.g., in standard equation modeling or use of random slopes as measures of individual change have also its strengths. IV) <u>Inclusion of covariates</u> representing other variables (e.g. neurochemical measurements) may account for unexplained variance and uncover further associations.

Here, we combined datasets to increase sample size and statistical power. However, we also inherited the idiosyncrasies of these datasets, such as inclusion/exclusion criteria, sample unrepresentativeness, and recruitment methods. Also, analytical compromises were made to ensure compatibility across all datasets. Memory function was harmonized independently within each dataset, and thus memory function reflects the specific tests used rather than a common construct. The Item-response framework is a more robust approach for harmonisation, placing all individuals in the same *space*; yet unfeasible in practice, as some datasets lacked shared tests. Estimating non-linear trajectories within individuals was also unfeasible given the relatively limited number of observations and follow-up durations per participant. The two-source model proposed in the discussion remains speculative, while the specific mechanisms underlying *brain aging* remain elusive. Young APOE ε4 carriers showed less memory change. We remain cautious of these results as the youngest segment of our sample had limited availability of APOE information, and thus emphasize the need for further research. Finally, this study is limited to regional cortical thickness and volume. Other macrostructural markers of brain aging such as neuromodulatory brainstem nuclei^83^, or even indices of brain function^84^, may provide additional insights.

### Conclusions

Regional brain decline over time, particularly, but not limited to, the hippocampus, was associated with memory loss with age, but APOE ε4 status was not a key factor behind these *change-change* associations. These associations strengthened with age from around 60 years. Methodologically, the findings underscore the necessity for methods and approaches that capture non-linear dynamics and put focus on the variability across individuals rather than mean change. Theoretically, the results support a multidimensional view of memory, aging and disease, where multiple factors converge to increase the vulnerability of common macrostructural systems supporting memory function in older age.

## Methods

### Participants

In this study we combined 13 ongoing or retrospective datasets that included a) cognitively healthy adult individuals with longitudinal assessments of brain structure (T1-weighted [T1w] sequence) and of memory function. All the main analyses were carried out using only longitudinal information from both brain structure and memory function. Individuals with only 1 observation, or uncoupled memory – brain data, were used only in preprocessing stages: for calibration purposes in MRI preprocessing and for principal component extraction, and Z-scoring of memory scores (see *below*). See **Supplementary Tables 5** and **6**, for information on the *initial MRI and memory samples.* Unless otherwise stated, we focus on the longitudinal-coupled samples used in the main analyses.

3,737 cognitively healthy adults, with – at least, partially – overlapping longitudinal follow-ups of brain structure and memory function, with a minimum total span of 1.5 years were included in the analyses. In total, 10,343 MRI observations and 13,460 memory observations contributed to the analyses (**Table 1, Supplementary Figure 1**). The datasets include the LCBC^85^, Betula^86^, UB^87,88^, and BASE-II^89,90^ datasets (from the Lifebrain Consortium)^91^ as well as the COGNORM^92^, the Alzheimer’s Disease Neuroimaging Initiative (ADNI) database (https://adni.loni.usc.edu)^93^, AIBL^94^, BBHI^95^, the Harvard Aging Brain Study (HABS)^96^, the UKB (https://www.ukbiobank.ac.uk/)^97^, PREVENT-AD^98,99^, OASIS3^100^, and VETSA^101^ datasets. In addition to cohort-specific inclusion and exclusion criteria, observations concurrent with cognitive impairment and Alzheimer’s dementia were excluded. Individuals with baseline age <18 years, or with severe neurological or psychiatric disorders were additionally excluded. Based on preprocessing requirements, MRI data from scanners with fewer than 25 observations were excluded as well as individuals with less than 1.5 years of follow-up either of memory function or brain structure. Individuals without partially overlapping follow-up periods of brain and memory assessment were excluded, as well as those with non-overlapping periods >10 years. See **Supplementary Table 7** for data availability, ethical standards, and contact information and **SI** for more sample details. A total of 3,149 subjects had APOE data available. Of these, 27.8 % were carriers of the APOE ε4 allele.

### Memory function

For each sample, we first z-normalized all measures based on the first time point and the different available memory tests. When multiple measures were available, we estimated a main component using Principal Component Analysis (PCA; *prcomp*) with all measures at the first time point as inputs. Missing values were imputed using *imputePCA* from the *missMDA* r-package^102^. Only for OASIS3, the imputed number of values was not negligible (> .5%). See **Supplementary table 8** for information of memory data for each sample. For each dataset, we regressed-out age as a smoothing term, sex, and one or two dummy test-retest regressors using GAMMs (*mgcv R-package*)^103^. Individual identifiers were used as random intercepts and the number of dummy test-retest regressors depended on whether the dataset had 2 or >=3 waves with memory function data. We retained individuals with at least 2 observations and a minimum follow-up of 1.5 years. For each individual, we then estimated yearly change by regressing memory observations on follow-up time, which were Z-standardized by site and fed to higher-level analyses.

### MRI preprocessing and brain structure

#### MRI acquisition and preprocessing

Structural T1w MPRAGE and FSPGR scans were collected using 1.5 and 3T MRI scanners. See information on scanner parameters and scanners across datasets in **Supplementary Table 9**. Data was converted to BIDS^104^ and preprocessed using the longitudinal FreeSurfer v.7.1.0 stream^105^ for cortical reconstruction and volumetric segmentation of the structural T1w scans^106,107^. See details in **SI.** Data was tabulated based on the *Destrieux* (cortical)^56^ and *aseg* (subcortical) atlases^57^.

#### Data harmonization

Brain regions were harmonized using a normative modelling framework resulting in site-agnostic deviation scores (*z*-scores) adjusted for age, and sex^59,79^ based on a Hierarchical Bayesian Regression technique^78^ as implemented in the *PCNtoolkit* (0.30.post2), in *Python3* environment^108^ (version 3.9.5). Calibration to the model was performed iteratively (N = 100) to avoid losing longitudinal observations. This step was carried out with the *initial MRI sample*, i.e. regardless of availability of longitudinal MRI data or paired memory function assessments. Calibrated data, across iterations, showed high reliability. See **SI** for more normative modelling harmonization details. Next, we selected individuals with at least 2 observations and a minimum follow-up of 1.5 years. For each individual and region, we estimated yearly change by regressing normative MRI values on follow-up time, Z-standardized data by site, and fed the output on higher-level analyses.

### Higher-level analyses

All the analyses were carried out in the R environment (version 4.2.1)^109^. Visualizations were made with the *ggplot2*^110^ and the *ggseg*^111^ R-packages. Analyses were mostly carried out using *gamm* models as implemented in the *mgcv R-package*^103^. Derivatives were estimated based on finite differences as implemented in the *gratia* package^112^. Linear mixed models as implemented in *lme4, lmerTest*^113,114^ were also used to assess the effect of APOE ε4 on brain and cognition.

To test the regional association between brain change and memory change, we carried out univariate weighted GAMMs, with a smooth term of Δbrain predicting Δmemory. An Δbrain × age tensor interaction term was added, as well as a smooth term of age, to assess the effect of age on Δbrain – Δmemory associations. The effect of APOE ε4 on memory and brain regions was tested using weighted linear mixed effects models, with APOE ε4 status predicting either Δbrain or Δmemory. The age × APOE ε4 status interaction was tested using GAMMs with age as a smooth term by APOE ε4 status as ordered factor, in addition to APOE ε4 status as fixed effect, and age as smooth term. This effectively models the smooth term of age for APOE ε4 non-carriers as reference while the smoothed term for APOE ε4 carriers models the difference with respect to the reference. Similarly, the Δbrain × APOE ε4 status was used to assess the effect of APOE ε4 on Δbrain – Δmemory associations. Finally, age × Δbrain × APOE ε4 status with their simple effects) was used to test a triple interaction of age, APOE ε4 status, and Δbrain on Δmemory. All models included dataset as random intercept. We tested the *dimensionality* by performing PCA and clustering on those regions (N = 19) showing significant Δbrain – Δmemory associations. The clustering was based on the M3C, Monte-Carlo Reference-based Consensus algorithm, implemented in the M3C package^115^ which, critically, tests whether the desired solution is better than K = 1. We used a *spectral clustering* algorithm and *PAC* criteria, while the remaining parameters were set to the default. As post-hoc analyses we tested whether the resulting clusters of brain change were related to memory change controlling for the effect on memory of other brain regions such as the hippocampus or a general factor of brain decline using GAMMs as described above. See more details on **SI** along with pseudocode.

Note that in all analyses we have one observation per individual (e.g. Δmemory) as we are using change scores. Note also that age (and sex) trends are removed and thus the model captures only interindividual associations - relative change - and age trends are uninterpretable. Prior to any analysis, outlier values, defined as values >4.5 SD from the mean, were removed from the analyses (based on a p < 0.05 of observing, at least, one outlier value given a normal distribution and our sample size). In GAMMs, we estimated p-values using a *wild* bootstrapping (n = 5000) as the out-of-the-box p-values, as implemented in *mgcv*, are anticonservative^81^. *Wild* bootstrapping generates a null distribution of p-values by a) estimating a *null* model without the regressor of interest, b) extracting predicted values from the model and its residuals, c) adding the predicted value to the residuals multiplied by a random vector of 1 and −1s, and d) re-estimating a new model using this score as your predicted variable. When appropriate, p-values were corrected for multiple comparisons using FDR^116^. All models used weights to account for unequal reliability of longitudinal data. That is, individuals with short follow-up periods and less observations contribute with more unreliable, high-variance data and thus should produce an unequal spread of residuals. We used the square of reliability as weights as estimated elsewhere^20^. Weights < .09, corresponding to longitudinal reliability < .3, were set at .09. For tensor interactions, we estimated the derivatives along Δbrain at specific ages (40, 50, 60, 70, 80 years) using a finite differences approach. The degree of association between Δbrain and Δmemory is estimated only in for Δbrain < 0 – as most associations are constrained in *brain decliners*, estimating the mean association across the Δbrain weighted by the density of data-points. Data from ventricles was sign reversed. We slightly trimmed the x-axis in the figures – removing ≈1 of the observations – to exclude high uncertainty fittings from visualization.

## Funding and Acknowledgements

This work was supported by the Department of Psychology, University of Oslo (to **K.B.W., A.M.F.**), the Norwegian Research Council (to **K.B.W.** [325001, 301395, 239889], **A.M.F.** [325878, 262453], **D.V.P** [324882]), the project has received funding from the European Research Council’s Starting Grant scheme under grant agreements 283634, 725025 (to **A.M.F.**) and 313440 (to **K.B.W.**), and the University of Oslo through the UiO:Life Science convergence environment [AHeadForLife: Societal and environmental determinants of brain and cognition] (to **A.M.F**). **R.N.H.** was supported by the UK Medical Research Council [SUAG/046/G101400]. A.P.-L. was partly supported by grants from the National Institutes of Health (R01AG076708), Jack Satter Foundation, and BrightFocus Foundation.

The different sub-studies are supported by different sources. **LCBC**: the Norwegian Research Council (to **A.M.F., K.B.W.**), and the National Association for Public Health’s dementia research program (**A.M.F.**). **Umeå (betula):** a scholar grant from the Knut and Alice Wallenberg (KAW) foundation to **L.N. UB**: **D.B.F.** was funded by ICREA Academia Award (2019) and 2014 awards from the Catalan Government. He acknowledges the CERCA Programme/Generalitat de Catalunya and is supported by María de Maeztu Unit of Excellence (Institute of Neurosciences, University of Barcelona) MDM-2017-0729, Ministry of Science, Innovation and Universities. **BASE-II (mpib).** BASE-II has been supported by the German Federal Ministry of Education and Research under grant nos 16SV5537, 16SV5837, 16SV5538, 16SV5536K, 01UW0808, 01UW0706, 01GL1716A and 01GL1716B and by the European Research Council under grant agreement no. 677804 (to **S.K.**). **BBHI.** The data from BBHI was obtained with funding from “la Caixa” Foundation (grant agreement n° LCF/PR/PR16/11110004), and also from Institut Guttmann and Fundació Abertis. **COGNORM** is funded by the South-Eastern Norway Regional Health Authorities (#2017095) The Norwegian Health Association (#19536) and by Wellcome Leap’s Dynamic Resilience Program (jointly funded by Temasek Trust) #104617). The funding sources had no role in the study design. Data used in preparation of this article were obtained from the **Alzheimer’s Disease Neuroimaging Initiative (ADNI) database** (adni.loni.usc.edu). The ADNI was launched in 2003 as a public-private partnership, led by Principal Investigator Michael W. Weiner, MD. The primary goal of ADNI has been to test whether serial magnetic resonance imaging (MRI), positron emission tomography (PET), other biological markers, and clinical and neuropsychological assessment can be combined to measure the progression of mild cognitive impairment (MCI) and early Alzheimer’s disease (AD). For up-to-date information, see https://adni.loni.usc.edu/. As such, the investigators within the ADNI contributed to the design and implementation of ADNI and/or provided data but did not participate in analysis or writing of this report. A complete listing of ADNI investigators can be found at: http://adni.loni.usc.edu/wp-content/uploads/how_to_apply/ADNI_Acknowledgement_List.pdf. Data collection and sharing for this project were funded by the ADNI (NIH Grant U01 AG024904). ADNI is funded by the National Institute on Aging, the National Institute of Biomedical Imaging and Bioengineering, and through generous contributions from the following: AbbVie, Alzheimer’s Association; Alzheimer’s Drug Discovery Foundation; Araclon Biotech; BioClinica, Inc.; Biogen; Bristol-Myers Squibb Company; CereSpir, Inc.; Cogstate Eisai Inc.; Elan Pharmaceuticals, Inc.; Eli Lilly and Company; EuroImmun; F. Hoffmann-La Roche Ltd and its affiliated company Genentech, Inc.; Fujirebio; GE Healthcare; IXICO Ltd.; Janssen Alzheimer Immunotherapy Research & Development, LLC.; Johnson & Johnson Pharmaceutical Research & Development LLC.; Lumosity; Lundbeck; Merck & Co., Inc.; Meso Scale Diagnostics, LLC.; NeuroRx Research; Neurotrack Technologies; Novartis Pharmaceuticals Corporation; Pfizer Inc.; Piramal Imaging; Servier; Takeda Pharmaceutical Company; and Transition Therapeutics. The Canadian Institutes of Health Research is providing funds to support ADNI clinical sites in Canada. Private sector contributions are facilitated by the Foundation for the National Institutes of Health (http://www.fnih.org). The grantee organization is the Northern California Institute for Research and Education, and the study is coordinated by the Alzheimer’s Therapeutic Research Institute at the University of Southern California. ADNI data are disseminated by the Laboratory for Neuro Imaging at the University of Southern California. Data used in the preparation of this article was obtained from the **Australian Imaging Biomarkers and Lifestyle flagship study of ageing (AIBL)** funded by the Commonwealth Scientific and Industrial Research Organisation (CSIRO) which was made available at the ADNI database (www.loni.usc.edu/ADNI). The AIBL researchers contributed data but did not participate in analysis or writing of this report. AIBL researchers are listed at www.aibl.csiro.au. Parts of the data used in the preparation of this article were obtained from the **Harvard Aging Brain Study** (**HABS** - P01AG036694; https://habs.mgh.harvard.edu). The HABS study was launched in 2010, funded by the National Institute on Aging. and is led by principal investigators Reisa A. Sperling MD and Keith A. Johnson MD at Massachusetts General Hospital/Harvard Medical School in Boston, MA.” **OASIS** data were provided [in part] by **OASIS 3**: Longitudinal Multimodal Neuroimaging: Principal Investigators: T. Benzinger, D. Marcus, J. Morris; NIH P30 AG066444, P50 AG00561, P30 NS09857781, P01 AG026276, P01 AG003991, R01 AG043434, UL1 TR000448, R01 EB009352. AV-45 doses were provided by Avid Radiopharmaceuticals, a wholly owned subsidiary of Eli Lilly. **PREVENT-AD** was funded by the Canadian Institutes of Health Research, McGill University, the Fonds de Recherche du Québec – Santé, Alzheimer’s Association, Brain Canada, the Government of Canada, the Canada Fund for Innovation, the Douglas Hospital Research Centre and Foundation, the Levesque Foundation, an unrestricted research grant from Pfizer Canada. Private sector contributions are facilitated by the Development Office of the McGill University Faculty of Medicine and by the Douglas Hospital Research Centre Foundation (http://www.douglas.qc.ca/). **UK Biobank** is generously supported by its founding funders the Wellcome Trust and UK Medical Research Council, as well as the Department of Health, Scottish Government, the Northwest Regional Development Agency, British Heart Foundation and Cancer Research UK. The organisation has over 150 dedicated members of staff, based in multiple locations across the UK. **VETSA** Parts of the data are from VETSA, which is funded by the National Institute of Aging grants R01s AG018384, AG018386, AG050595, AG022381, AG076838. The content is the responsibility of the authors and does not necessarily represent official views of the NIA, NIH, or VA. U.S. Department of Veterans Affairs, Department of Defense; National Personnel Records Center, National Archives and Records Administration; Internal Revenue Service; National Opinion Research Center; National Research Council, National Academy of Sciences; and the Institute for Survey Research, Temple University provided invaluable assistance in the conduct of the VET Registry. The Cooperative Studies Program of the U.S. Department of Veterans Affairs provided financial support for development and maintenance of the Vietnam Era Twin Registry. We would also like to acknowledge the continued cooperation and participation of the members of the VET Registry and their families.

## Author Contributions

**D.V.P.** Conceptualization, Methodology, Formal analysis, Writing - Original Draft; **Ø.S.** Methodology, Software, Formal analysis, Writing - Review & Editing; **M.S.** Resources, Data Curation, Writing - Review & Editing; **W.B.** Writing - Review & Editing; **I.K.A.** Software, Writing - Review & Editing; **D.B-F.** Resources, Writing - Review & Editing; **A.B.** Writing - Review & Editing; **G.C.** Resources, Writing - Review & Editing; **S.D.** Resources, Data Curation, Writing - Review & Editing; **P.G.** Methodology, Writing - Review & Editing; **R.N.H.** Conceptualization, Writing - Review & Editing; **S.K.** Writing - Review & Editing; **U.L.** Resources, Writing - Review & Editing; **A.M.M.** Software, Writing - Review & Editing; **L.N.** Conceptualization, Resources, Writing - Review & Editing; **A.P.L.** Conceptualization, Resources, Writing - Review & Editing; **J.M.R.** Data Curation, Writing - Review & Editing; **J.S.S.** Resources, Writing - Review & Editing; **C.S.P.** Resources, Writing - Review & Editing; **L.O.W.** Resources, Writing - Review & Editing; **T.W.** Methodology, Software, Formal analysis, Writing - Review & Editing; **K.B.W.** Conceptualization, Resources, Writing - Review & Editing; **A.M.F.** Conceptualization, Resources, Writing - Review & Editing.

## Materials & Correspondence

Correspondence should be addressed to Didac Vidal Pineiro.

## Data availability

The raw data were gathered from 13 different datasets. Different agreements are required for each dataset. Most dataset are openly available with prespecified data usage agreements. For some datasets, such as UKB, fees may apply. Requests for Lifebrain cohorts (LCBC, Umeå, UB) and COGNORM, should be submitted to the corresponding principal investigator. See data availability and contact details for all datasets in **Supplementary Table 7.**

## Code Availability

Statistical analyses in this manuscript are available alongside the manuscript and will be made available at https://github.com/daidak/memory-brain-change. All analyses were performed in R. The scripts were run on the Colossus processing cluster, University of Oslo. MRI preprocessing and feature generation scripts were performed with FreeSurfer (https://surfer.nmr.mgh.harvard.edu/) software.

