## Supplementary Information for "Vulnerability to memory decline in aging – a mega-analysis of structural brain change"

### Methods

#### Participants

In this study we combined 13 ongoing or retrospective datasets that included a) cognitively healthy adult individuals with longitudinal assessments of brain structure (T1-weighted scans) and memory function. All the main analyses were carried out using only longitudinal data from both brain structure and memory function. Individuals with only one observation, or uncoupled memory – brain data, were used only in preprocessing stages: for calibration purposes in MRI preprocessing and for principal component extraction, and Z-scoring of memory scores (see *below*). See **Supplementary Tables 5** and **6,** for information on the *initial MRI and memory samples.* Unless otherwise stated, we will focus on the longitudinal-coupled samples used in the main analyses.

3737 unique cognitively healthy adults, with – at least partially – overlapping longitudinal follow-ups of brain structure and memory function, with a minimum total span of 1.5 years were included in the analyses. In total, 10343 MRI observations and 13460 memory observations contributed to the analyses (see **Table 1**). The datasets include the LCBC[1], Betula[2], UB[3,4], and BASE-II[5,6] datasets (from the Lifebrain Consortium)[7] as well as the COGNORM[8], the Alzheimer's Disease Neuroimaging Initiative (ADNI) database (https://adni.loni.usc.edu)[9], AIBL[10], BBHI[11], the Harvard Aging Brain Study (HABS)[12], the UKB (https://www.ukbiobank.ac.uk/)[13], PREVENT-AD[14,15], OASIS3[16], and VETSA[17] datasets. In addition to cohort-specific inclusion and exclusion criteria, observations concurrent with cognitive impairment and Alzheimer's dementia were excluded. Individual with baseline age <20 or with severe neurological or psychiatric disorders were additionally excluded.

Based on preprocessing requirements, MRI data from scanners with fewer than 25 observations were also excluded based on stability issues in the normative modeling-based harmonization. For computing slopes, individuals with less than 1.5 years of follow-up, either of memory function or brain structure were also excluded. For memory–brain pairing, individuals without at least partially overlapping follow-up intervals for brain structure and memory were excluded, as well as those in which one of the assessments (in practice, memory function only) started > 10 years before or ended > 10 years after the beginning or end of the

MRI follow-up period. The initial dataset included individuals with 1 to 14 MRI acquisitions with longitudinal structural MRI data spanning up to 15.8 years. Similarly, memory assessments range from 1 to 30 observations per individual with a follow-up up to 31.6 years. See **Supplementary Table 7** for data availability, ethical standards, and contact information.

ADNI: The Alzheimer's Disease Neuroimaging Initiative (ADNI)[9] is a multi-site project led by Doctor Michael W. Weiner to assess the progression of mild cognitive impairment (MCI) and early Alzheimer's Disease (AD), combining imaging, clinical and other biological markers, and neuropsychological and clinical assessments over time. For more information, visit https://adni.loni.usc.edu/about/. The age range for the participants is 55-90 years. The present study includes participants from ADNI 1, ADNIGO, ADNI2, and ADNI 3, who were cognitively healthy at baseline (*DX_bl* variable). Only observations in which participants were still cognitively healthy were included as determined by the ADNI team (*DX* variable). Amongst others, participants were required to have no evidence of ischemic stroke (Hachinski Ischemic Score ≤ 4), a Geriatric Depression scale score < 6, stable medications for 4 weeks before the screening, good auditory and visual acuity, good general health, no medical contraindications to MRI and at least 6 grades of education/work history. In-detailed general inclusion and exclusion criteria are described elsewhere[18]. All participants signed an informed consent form, and the protocols were approved by the corresponding regional ethical committees in the US and Canada. Data was retrieved in April 2021.

AIBL: The Australian Imaging, Biomarker & Lifestyle Flagship Study of Ageing (AIBL)[10] is a prospective study including cognitively normal participants, and patients with mild cognitive impairment, or AD aged 60 years or older. The study assesses the biomarkers, genetic factors, cognitive characteristics, and health and lifestyle factors that are associated with the development of AD combining techniques such as MRI, positron emission tomography (PET), blood tests, and fluid sample analysis, as well as neuropsychological and clinical assessments. Healthy participants must meet specific criteria, including being free of cognitive impairments and having test performance within 1.5 SD of age-adjusted norms. The test battery and sample description as well as in-detail general inclusion criteria have been described in detail previously[10]. All observations corresponding to cognitively normal individuals were included, as defined and evaluated by the AIBL team. All participants signed an informed consent form and the protocol was approved by the institutional human research committees of Austin

Health, St Vincent's health, Hollywood Private Hospital, and Edit Cowan University (Australia). Data was retrieved in February 2023.

BASE-II: Participants of the Berlin Aging Study II (BASE-II; mpib) were community-dwelling older adults recruited from the greater Berlin metropolitan area through advertisements in newspapers and public areas[5,6]. The baseline sample comprised 2200 participants; 1600 older adults aged 61–88 years, and 600 younger adults aged 24–40 years. Participants were invited to a medical exam and cognitive testing sessions. After completion of the cognitive examination of BASE-II, eligible participants were invited to take part in one MRI session within a time window of 2–4 weeks after cognitive testing. The MRI sample consisted of 341 older adults aged 61–82 years and 103 younger adults. MR scans and cognitive scores were obtained 2012-2013. A subsample of the MR sample was later re-invited for follow-up. The different elements of the study were approved by the ethics committees of the Max Planck Institute for Human Development, the Charité University ethics committee and by the ethics committees of The German Association for Psychology (DGPs). Participants signed written informed consent and received monetary compensation for their participation in BASE-II and the MRI study. All experiments were performed in accordance with relevant guidelines and regulations. Inclusion criteria for taking part in this study were age between 20 and 35 or 60 and 80 years, apparently healthy. Exclusion criteria were untreated diabetes and hypertension; prior stroke, head injuries or brain surgery; psychiatric illness; major depression; dementia with a score < 24 on the Mini- Mental State Examination. None of the participants took medication that might affect memory function or had a history of head injuries, medical (e.g., heart attack), neurological (e.g., epilepsy), or psychiatric disorders (e.g., depression). The BASE-II cohort was part of the Lifebrain obtained as part of the Lifebrain consortium[7]. All participants reported normal or corrected to normal vision. Observations with MMSE < 26 or no MMSE data were discarded. Data was retrieved in February 2022.

BBHI: Barcelona Brain Health Initiative study (BBHI; https://bbhi.cat/en/) participants are community-dwelling individuals between 40 and 65 years of age, of both sexes, free from any self-reported neurological or psychiatric diagnosis at the time of recruitment[11,19]. It is an ongoing longitudinal cohort study that aims investigating the determinants of brain and mental health in healthy middle-aged and older adults. Recruitment started in 2017, when multiple initiatives (including conferences, radio and television interviews, and social media advertisements) took place to encourage participants to join the study. It has enrolled 4,686 participants via a web-based application which completed a first on–line questionnaire. The BBHI includes periodic cognitive, medical, brain imaging, and biological assessments. BBHI has several sub studies. In one of them, a sample of 1,000 participants is undergoing a detailed clinical phenotyping through a multi-day in-person evaluation that includes cognitive, physical, and medical assessments, biological sample recollection, structural and functional magnetic resonance imaging (MRI), and electroencephalography (EEG). Participants of this study are invited biannually for repeated evaluations. Data was retrieved in July 2023.

BETULA: The BETULA project (Umeå)[20,21] is a prospective longitudinal study on aging, memory, and dementia, which used a population-based sampling of healthy middle-aged and older adults for recruitment. Detailed recruitment procedures are found elsewhere[20,21]. For the current analyses, the MRI subsample of the study is used. Participation in the neuroimaging study was offered to all participants who had remained in the study and completed cognitive testing at the 5th Betula test wave onwards. Exclusion criteria were severe visual or auditory handicaps, intellectual or developmental disabilities, suspected dementia, having a mother tongue other than Swedish, MRI contraindications, severe neurological disorders, or visual/motor deficits that could interfere with fMRI data collection, MMSE <24, brain or head surgery, and substantial brain anatomical deviations. Some participants were later excluded due to discovered neurological conditions, severe depression, and MRI anatomical abnormalities. All participants signed an informed consent, and the protocols were approved by the Regional Ethical Vetting Board at Umeå University. The BETULA cohort was part of the Lifebrain obtained as part of the Lifebrain consortium[7]. Data was retrieved in August 2022.

COGNORM: The COGNORM cohort[22] is an ongoing, prospective study coordinated by the Oslo University Hospital and Diakonhjemmet Hospital, Oslo, Norway. Patients (age ≥ 65 years) scheduled for elective gynecological, urological, or orthopedic surgery under spinal anesthesia were recruited. Participants were required to have no dementia, previous stroke with sequela, Parkinson’s disease, or other neurodegenerative diseases that are likely to affect cognition. Patients with suspected undiagnosed dementia at any time within the first five years of follow-up (n = 15)[23], MMSE score <28 at baseline, and at least two abnormal cognitive test scores (-1.5 standard deviation [SD] below the mean normal value for age, sex, and education) were excluded. All observations corresponding to cognitively normal individuals were included. All participants signed an informed consent form, and the protocol was approved by the Norwegian Regional Committees for Medical and Health Research Ethics and the Data Protector Officer at Oslo University Hospital. Data was retrieved in March 2023.

HABS: The Harvard Aging Brain Study (HABS)[24] is an ongoing, long-term observational study that aims to enhance our understanding of brain aging and the early stages of Alzheimer's disease. The study collects PET, MRI data, neuropsychological and clinical assessments. The age range was between 50 and 90 years at the time of baseline assessment, and all patients were considered non-clinically impaired at the start of the study. Further participants had a CDR score of 0, MMSE score ≥ 25, < 11 on the Geriatric Depression Scale, and scores above age- and education-adjusted cutoffs on the 30-Minute Delayed Recall of the Logical Memory Story A to be included in the study. Participants with a history of alcoholism, drug abuse, head trauma, or current serious medical/psychiatric illness were excluded. Further details can be found elsewhere[24]. Observations with MCI or AD diagnostic (DX variable) were excluded. All participants signed an informed consent form, and the protocol was approved by the Partners Healthcare Human Research Committee. Data consisted of the "HABS data release 2.20", retrieved in August 2022 via habs.mgh.harvard.edu.

LCBC: The Center for Lifespan Changes in Brain and Cognition cohort (LCBC, Oslo)[1,25] consists of cognitively healthy, community-dwelling participants across the lifespan and is drawn from studies coordinated by the LCBC Research Group (LCBC www.oslobrains.no), approved by a Norwegian Regional Committee for Medical and Health Research Ethics. Written informed consent was obtained from all participants. The samples were recruited by a variety of methods such as newspapers and webpage ads. Most participants were recruited for observational studies, some currently ongoing, while a minority were recruited to enter into cognitive training. Written informed consent was obtained from all adult participants. All participants had to undergo a standardized health interview before being included in the study, and those with a history of neurological or psychiatric conditions or who reported concerns about their cognitive function were excluded. Additionally, all participants over the age of 40 years were required to score at least 25 on the Mini-Mental State Examination. The LCBC cohort was part of the Lifebrain obtained as part of the Lifebrain consortium[7]. MRI observations paired with MMSE ≤ 25 were excluded. Data was retrieved in November 2022.

OASIS3: The Open Access Series of Imaging Studies (OASIS3)[26] is a retrospective collection of multimodal data that focuses on aging and AD and is openly accessible to the scientific community. OASIS-3 includes neuroimaging, clinical and neuropsychological data. Participants were recruited through the Washington University Knight Alzheimer Disease Research Center via flyers, word of mouth, and community engagements and were aged between 42 and 95 years. Only participants deemed cognitively normal at baseline were included in the observations. Exclusion criteria included medical conditions that precluded longitudinal participation or medical contraindications for the different study arms. See in-detail inclusion and exclusion criteria[26]. All participants consented to Knight ADRC-related projects following procedures approved by the Institutional Review Board of Washington University School of Medicine. Observations were included until the last observation in which a subject was deemed cognitively healthy as determined by the Clinical Dementia Rating Scale (CDR)[27]. Data was retrieved in November 2022.

PreventAD: The Pre-symptomatic Evaluation of Experimental or Novel Treatments for AD (PREVENT-AD)[28] is a retrospective, long-term study that follows cognitively healthy older individuals with a familiar history of AD. It includes participants enrolled either from an observational cohort or the clinical trial of PREVENT-AD. This study comprises MRI images, blood and CSF samples, and clinical and neuropsychological assessments. Participants in the study had to be at least 60 years old, had ≥6 years of education, and they needed to be cognitively unimpaired at baseline. The Montreal Cognitive Assessment (MoCA) and CDR scales were used to assess cognitive abilities, and participants were considered cognitively intact if their MoCA scores were ≥ 26/30 or their CDR was = 0. Other exclusion criteria at baseline included medical conditions that prevented longitudinal participation or medical contraindications to MRI, use of acetylcholinesterase inhibitors, other approved prescription cognitive enhancers, hypertension, or substance abuse. The inclusion and exclusion criteria have been previously described in detail[28]. The protocols, consent forms, and study procedures were approved by the McGill Institutional Review Board and the Douglas Mental Health University Institute Research Ethics Board. Observations with RBANS > 1SD below the mean and probable MCI, as evaluated by a clinician, were excluded. Data was retrieved in February 2022.

UB: The University of Barcelona cohort consisted of a series of retrospective sub studies[3,4,29,30]. In all cases, samples consisted of cognitively healthy, community-dwelling participants with normal cognitive and visual function. Most participants were recruited for observational studies, while a minority were recruited to enter into cognitive training. Exclusion criteria varied a bit across studies (see specific studies for details), but include severe neurologic and psychiatric disorders, recent head trauma or brain surgery, cognitive deterioration, or dementia with a score < 24 on the Mini-Mental State Examination and additional neuropsychological criteria, other neurodegenerative disorders like Parkinson's disease and chronic illness with a projected shortened lifespan. Follow-up observations with MMSE < 26 were excluded. All participants signed an informed consent, and the protocols were approved by the ethical committees of the University of Barcelona and of the Hospital Clinic of Barcelona. The UB cohort was part of the Lifebrain obtained as part of the Lifebrain consortium[7]. Data was retrieved in February 2022.

UKB: The UK Biobank (UKB) (https://www.ukbiobank.ac.uk/about-biobank-uk/) is a major national and international health resource with the aim of improving the prevention, diagnosis and treatment of a wide range of illnesses. UK Biobank recruited ≈500,000 people aged between 40-69 years in 2006-2010 from across the country to take part in this project[31]. Potential participants were identified through National Health Service (NHS) registers according to being aged 40-69 and living within a reasonable traveling distance of an assessment center. Assessment centers are located in accessible and convenient locations with a large surrounding population. Participants have undergone measures and provided samples and detailed information about themselves and agreed to have their health followed. The study sample was drawn from the UK Biobank neuroimaging branch[13] and conducted under data application number 32048. Only individuals with longitudinal MRI data were used in this study. Note that MRI showcase data was not used. Rather, MRI data was re-processed inhouse with the longitudinal pipeline. Participants signed an informed consent and the protocols were approved by the North West Multi-Center Research Ethics Committee [MREC]; see also https://www.ukbiobank.ac.uk/the-ethics-and-governance-council.

VETSA: The Vietnam Era Twin Study of Aging (VETSA) is an ongoing large-scale investigation of cognitive aging and brain aging in men[17,32,33]. It primarily investigates genetic and environmental influences on cognitive aging, brain structure and function, and health. VETSA involves over 1600 male twins from the Vietnam Era Twin Registry who served during the Vietnam War era. Assessments began when participants were in their 50s (in 2003) and follow-ups are conducted every 5-6 years. The age range of participants is narrow (from 52 to 60 at baseline). It has, amongst other, extensive neurocognitive testing, legacy cognitive data, genome, MRI, and plasma biodata. Over 1200 twins participated in waves 1, 2 and 3. All VETSA participants are men and all were in some branch of military service at some time between 1965 and 1975, though approximately 80% report no combat experience. The sample is a reasonably representative, community-dwelling sample of men in their age range in USA. Attrition-replacement procedures were taken in wave 2. VET registry was created by inviting to participate male twins who served in the United States military during the Vietnam Era by identifying them through their birth year (between 1939–1957), period of active duty (between 1965–1975), and having the same last name, different first name, same date of birth, and same first five digits of the Social Security number. VETSA twins are recruited from a participant pool (n = 3322) of a previous research study (Harvard Twin Study of Drug Abuse and Dependence) which attempted to contact all available VET Registry members (n = 7375). Recruitment for the VETSA study involved identifying eligible twins through the VET Registry at the VA Medical Center in Seattle. A survey firm then contacted potential participants with an introductory letter, followed by a phone call. Interested individuals submitted an authorization form indicating consent and site preference. These forms were processed and forwarded to the respective study sites, which then contacted participants. For the VETSA MRI study, participants were screened for safety issues (e.g. MRI contraindications), and both members of a twin pair had to consent to participate. For MRI, twins had to additionally (be able) to travel to a scanning site. Other exclusion criteria depended on exclusion criteria for serving in the military, e.g., participants scoring in the lowest 10 percentile ranks of the Armed Forced Qualification Test (AFQT) were excluded from the military. In addition, we further excluded data from participants with incidental radiological findings, history of seizure, and diagnostic of multiple sclerosis, and AIDS. Data was retrieved in July 2023.

#### MRI Acquisition, preprocessing, and harmonization

**MRI acquisition and preprocessing.** Structural T1-weighted (T1w) MPRAGE and FSPGR scans were collected using 1.5 and 3T MRI scanners. See information on scanner parameters and scanners across datasets in **Supplementary Table 9**. For datasets not provided in Brain Imaging Data Structure (BIDS) format, data was converted to BIDS[34]. BIDS transformation of ADNI, AIBL, OASIS3, and HABS data was performed with Clinica software[35,36]. We used the longitudinal FreeSurfer v.7.1.0 stream[37] for cortical reconstruction and volumetric segmentation of the structural T1w scans[38,39]. For sessions with multiple scans, data from the scanners were averaged. Briefly, the images were processed using the cross-sectional stream, which includes the removal of nonbrain tissues, Talairach transformation, intensity correction, tissue and volumetric segmentation, cortical surface reconstruction, and cortical parcellation. Next, an unbiased within-subject template space based on all cross-sectional images was created for each participant, using robust, inverse-consistent registration[40]. The processing of each time point was then reinitialized with common information from the within-subject template to increase reliability and statistical power. Except for the BETULA dataset, all data was preprocessed on the Colossus processing cluster, part of the Tjenester for Sensitive Data (TSD) (https://www.uio.no/tjenester/it/forskning/sensitiv/), University of Oslo (UiO). Data was tabulated based on the *Destrieux* (cortical)[41] and *aseg* (subcortical) atlases[42].

**Data harmonization.** Brain regions were harmonized using a normative modelling framework[43,44] with the *PCNtoolkit* (0.30.post2), in *Python3* environment[45] (version 3.9.5). This framework offers several advantages as it i) is carried independently across sites, ii) can isolate site-effects from other sources of variance associated with it, and iii) produces site-agnostic deviation scores (*z*-statistics) adjusted for age and sex. *PCNtoolkit* uses a Hierarchical Bayesian Regression (HBR) technique[46] and pretrained models from 82 different datasets, including the UKB. New sites can be added to the model using a held-out calibration dataset. To avoid losing longitudinal observations, here, we performed calibration recursively by iteratively (n = 100) holding out a calibrating sample and computing the estimates on the remaining data. The average scores of all iterations were used as the standardized scores for each observation. Scanners contributing with < 12 unique individuals or < 25 observations were removed. For scanners contributing > 12 and < 32 unique individuals, we used a calibration sample consisting of all but 2 participants and estimated the harmonized scores in these two. For scanners with >= 32 unique individuals, we used, in each iteration, a held-out sample of 30 individuals while estimates were applied to the rest. Most scanners with < 32 unique individuals belonged to ADNI due to its multisite nature. Note this step was performed with the *initial MRI sample*, i.e., regardless availability of longitudinal MRI data or paired memory function assessments. In most regions, normative scores show high reliability across iterations (intraclass correlation across regions [ICC] (.97[.01], range = .95 - .97). For longitudinal brain change, the mean reliability was even higher (.98[.01], range = .97 - .98). This is expected, since the uncertainty (error) is shared across observations of the same individual as long as the scanner does not change, i.e., the most recurring scenario. The correlation between harmonization with a normative modelling framework and Generalized Additive Mixed Models (GAMMs) – using age as a smoothing term, sex as a fixed factor, and dataset and scanner as random intercepts was r = 0.98 (0.03). The only regions with a correlation r < 0.90 were ventricle regions. This is likely due to increased dispersion of ventricular volume with age. Normative modelling models age variations in variability and converts the data to Z-scores based on age, and sex, while GAMMs model age changes in the mean but not in the variability. Next, we selected individuals with at least 2 observations and a minimum follow-up of 1.5 years. The follow-up criterion were chosen to exclude data with very low reliability for capturing *long-term* brain change, i.e., brain aging[47]. For each individual and region, we estimated yearly change by regressing normative MRI values on follow-up time (Δbrain). Brain change datawas Z-standardized by each site and coupled with memory change data for higher-level analyses. In practice, Δbrain reflects the yearly brain change of an individual relative to their age and sex peers. For pairing Δbrain with Δmemory, individuals had to have – at least partially - overlapping follow-up intervals, while the non-overlapping period had to be lower than 10 years.

### Results

#### Hemispheric asymmetries in the association between brain change and memory change.

No strong evidence for asymmetry in change – change associations was found (paired t-test; $\Delta\beta_w$[left-right] = 0.007, $t(82) = 1.78$, $p = 0.079$) (**Supplementary Figure 2a**). However, leftward asymmetry in change – change association was found at age 50 ($t(82) = 2.66$, $p_{FDR} = 0.05$). Asymmetry estimates in change – change associations decrease with increasing aging ($\Delta\beta_w$[left-right] = 0.017, 0.016, 0.008, 0.001, -0.002 at 40, 50, 60, 70, and 80 years of age) (**Supplementary Figure 2b)**.

#### Independent regional contributions to memory decline.

We employed GAMMs to investigate whether brain changes within specific clusters (refer to **Figure 3, Supplementary Table 3**) were associated with memory change. We conducted 4 analyses: i) basic, without additional covariates; ii) controlling for the hippocampus-based cluster; iii) controlling by the main factor (PC1) of brain decline; and iv) including all clusters together in a single model. Dataset was used as a random intercept. Detailed statistics are available in **Supplementary Table 4**. All clusters showed significant Δbrain – Δmemory associations without adjusting for covariates (all p's ≤ .001). This aligns with expectations, since all regions in the clustering analyses demonstrated associations with memory change. When controlling for the hippocampal cluster (#1), five clusters showed significant Δbrain – Δmemory associations: cluster #3, the left and right amydala ($\beta_w$ = .11, p = 0.01); cluster #5, left pericallosal sulcus, right gyrus rectus, and right subcallosal gyrus ($\beta_w$ = .16, p = 0.002), cluster #6, right long insular gyrus and planum polare ($\beta_w$ = .06, p = 0.01); cluster #7, right temporal transverse sulcus ($\beta_w$ = .05, p = 0.02); and cluster #8, left parahippocampal gyrus ($\beta_w$ = .05, p = .03). When controlling for a main factor of global decline, four clusters showed significant Δbrain – Δmemory associations: cluster #3, the left and right amydala ($\beta_w$ = .09, p = 0.05); cluster #5, left pericallosal sulcus, right gyrus rectus, and right subcallosal gyrus ($\beta_w$ = .11, p = 0.01), cluster #7, right temporal transverse sulcus ($\beta_w$ = .03,p = 0.01); and cluster #4, right supramarginal gyrus ($\beta_w$ = -.07, p = .02). The lack of Δbrain – Δmemory associations for the hippocampal cluster is probably explained by the fact that the factor of global decline weights heavily on the left and right hippocampus (**Figure 3c**). When included in a single model, five clusters showed significant Δbrain – Δmemory associations: cluster #1, the left and right hippocampus and left inferior lateral ventricle ($\beta_w$ = .09, p = 0.001); cluster #3, the left and right amydala ($\beta_w$ = .08, p = 0.03); cluster #5, left pericallosal sulcus, right gyrus rectus, and right subcallosal gyrus ($\beta_w$ = .11, p = 0.01), cluster #7, right temporal transverse sulcus ($\beta_w$ = .03,p = 0.04); and cluster #8, left parahippocampal gyrus ($\beta_w$ = .07, p = .02). Note that $\beta_w$ was estimated on above-average brain decliners, due to the non-linear nature of the associations (**Figure 1**). Note also that all p-values, except for the single model analysis, were obtained via bootstrapping. For the single model analysis, the p-values were derived from the implementation in the *mgcv R-package*[48] as bootstrapping proved overly conservative, possibly due to cross-correlations between clusters.

#### Replication using APOE ε4 non-carriers-only dataset.

To evaluate the potential influence of APOE ε4 on the Δbrain – Δmemory associations, we conducted analyses exclusively within a sample of APOE ε4 non-carriers and compared the results to the overall sample. 99.94% of the estimates in APOE ε4 non-carriers fell within the confidence interval of the estimates from the overall sample. 17 out of 19 regions that were significant in the entire sample were also significant ($p < .05$) in the APOE ε4 non-carrier sample. The exceptions were the left parahippocampal and right supramarginal gyri, where Δbrain – Δmemory associations did not reach significance ($p = 0.05$). Further, the influence of age on the Δbrain - Δmemory association was also consistent within the APOE ε4 non-carrier sample. 92.18% of the estimates from APOE ε4 non-carriers fell within the confidence intervals of the main sample. The left putamen, right parahippocampal gyrus, and left temporal pole showed the least consistency between both samples. Approximately, 91% of the estimates that fell outside the main sample's confidence intervals pertained to estimates in the younger age range (age ≤ 50). All 7 regions that showed a significant effect of age on Δbrain–Δmemory associations were significant in the APOE ε4 non-carrier sample.

#### Mechanisms behind change – change associations. A post-hoc simulation analysis.

We performed a simulation to explore potential explanations for our empirical findings, particularly the non-linear *change – change* associations, the moderating effect of age, and the lack of APOE ε4 effects. Using the *sn* package[49], we developed a simplified schematic model where the *observed* brain data resulted from two underlying sources. The first source represented *brain aging (true* or *latent* brain change)*,* characterized by a negatively skewed distribution with mean decline indicative of long-term, *degenerative* changes. The second source was modeled as a Gaussian distribution centered around zero, representing measurement error and other short-term influences[47,50,51]. Memory decline was modeled as linearly related to the *brain aging* component, plus random Gaussian noise. To explore the moderating effects (or lack thereof) of age and APOE ε4, we adjusted the parameters of the *brain aging* component, including mean, dispersion (e.g. variability), and skewness.

We generated independent *brain aging* and *measurement error* distributions (N = 3700) with the *rsn* function (*sn r-package*). This function is parametrized by location (ξ), scale (ω), and shape (α), which corresponds to the distribution's central tendency, dispersion, and skewness, respectively. Note that ξ and ω equate to mean (μ) and SD (σ) only when α = 0. The *measurement error* distribution was parametrized with ξ = 0, ω = .8, and α = 0; i.e., a Gaussian distribution with μ = 0 and σ = .8. The *main* brain aging was defined with ξ = -.8, ω = 1.2, and α = -5; producing a distribution characterized by a mean decline, a standard deviation comparable to that of the measurement error, and a negative (left) skew.

Most individuals exhibited values < 0, with a few showing accelerated decline (more negative values). Memory decline was modeled as linearly related to the *brain aging* component (r = .3), plus random Gaussian noise (ξ = 0, ω = .8). To test the effect of the mean, and dispersion on the *change – change* associations, we created distributions with ξ = -.2, -.8, and -1.2 and ω = .6, 1.2, 1.8, respectively. To test the effect of skewness, while maintaining variance ($\sigma^2$) constant, we used α = -2, -5, -20 and adjusted ω accordingly to maintain consistent dispersion. We ran 1000 iterations for each model, using GAMs to fit Δbrain – Δmemory associations. The mean trajectories and their 95% confidence intervals are presented in **Figure 7**.

The parameters selected for our simulations are somewhat arbitrary, as they can vary significantly across different regions and populations, e.g. younger vs. older individuals[47]. However, they are roughly comparable to those typically observed in studies of regional brain changes in cognitively healthy older adults. Focusing on hippocampal atrophy, previous research has reported annual hippocampal volume loss rates of -0.84%[52] and -1.05%[47], aligning with the $\xi$ = 0.8% used in our simulations. Assessing variability is more complex, as the results depend on the relationship between variability in latent brain change (brain aging) and measurement noise, which here corresponds to uncertainty in the estimated slopes of brain change. This ratio is influenced by study duration and the number of observations[52]. In prior work, we reported a standard deviation ($\sigma$) of 0.6 for yearly hippocampal atrophy rates - compared to $\sigma \approx 0.74$ used in our simulations - and a $\sigma$ of 1.23 for cross-sectional measurement error. The ratio used in the simulations would roughly match a study design involving three observations over approximately 3 to 3.5 years. Information on skewness is scarce, though we reported mild negative skewness in the distribution of observed brain change[47]. Assuming Gaussian noise, the latent brain change distribution would exhibit more pronounced negative skewness. The degree of skewness attenuation from *latent* to *observed* brain change also depends on the variability of the noise. In our simulated data, the *brain aging* (*latent*) skewness is $\gamma_1 = -0.85$, while the *observed* skewness is $\gamma_1 = -0.28$, which is entirely consistent with our previous estimations. Further research is needed to comprehensively characterize the distributions of brain changes.

### Figures and Tables

#### Supplementary Figure 1. Age distribution of the different datasets.

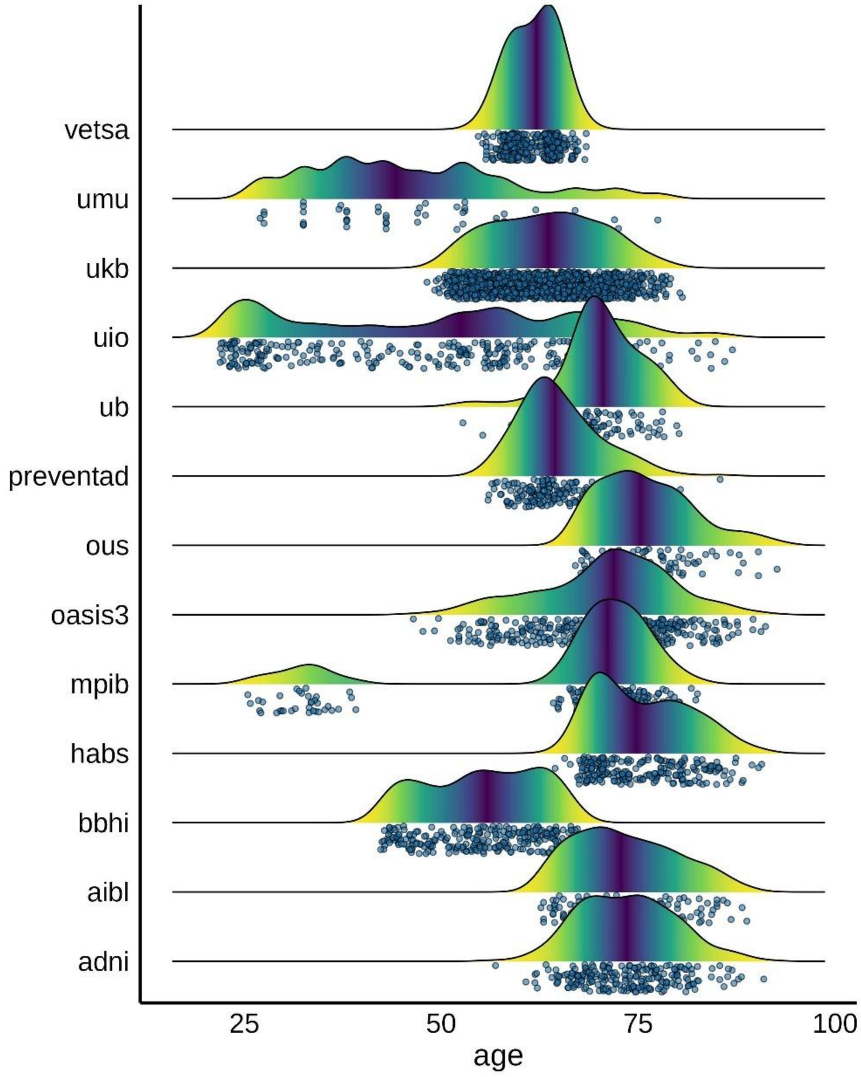


***Supplementary Figure 1. Age distributions.*** *Age distributions for the different datasets. Each point represents one individual at its mean age.*

#### Supplementary Figure 2. Hemispheric asymmetry in Memory change – brain change associations

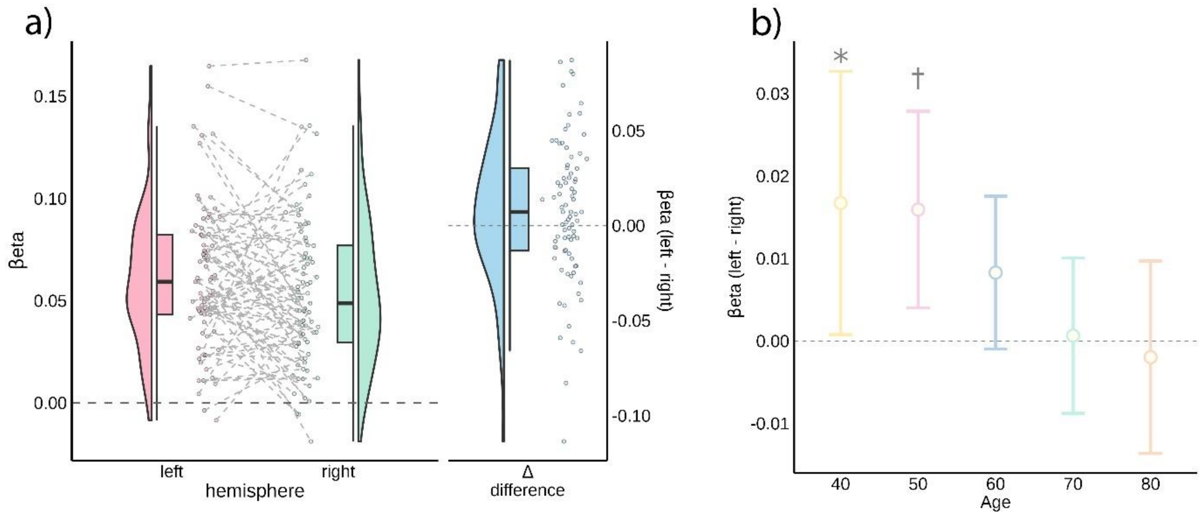


***Supplementary Figure 2. Memory change – brain change associations. Hemispheric asymmetry. a)*** *Left panel: Estimates ($\beta_w$) for Δbrain - Δmemory associations in the left and right hemispheres per regions. Right panel hemispheric differences in estimates between left and right hemispheres. Estimates were estimated as the density-weighted mean correlation between Δbrain and Δmemory, using the derivative function. Each point represents a region.* ***b)*** *Mean hemispheric difference and 95% CIs in regional estimates of Δbrain and Δmemory associations as a function of age. * denotes uncorrected $p < .05$; † denotes corrected $p_{FDR} < .05$.*

#### Supplementary Figure 3. Consensus Clustering results

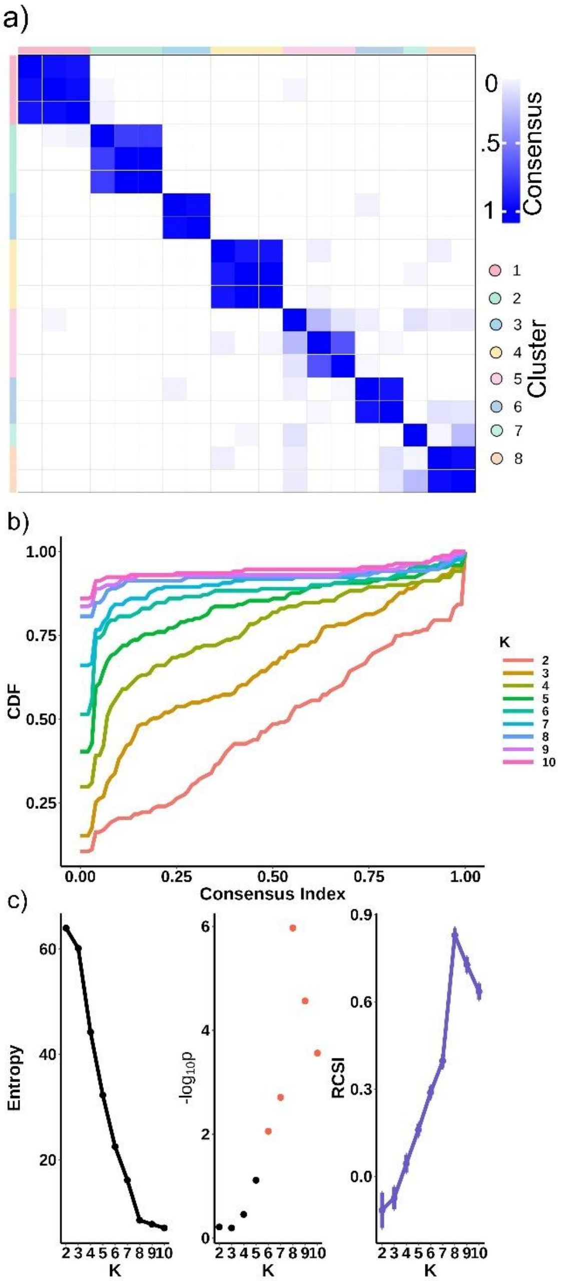


***Supplementary Figure 3. Performance metrics of consensus clustering.*** *a) Consensus clustering matrix at the selected solution (K = 8). The white-blue scale represents the proportion of iterations two regions have been grouped together in the same cluster. Ideally, values for between-cluster regions are 0, and for within-cluster regions are 1. b) Cumulative density function (CDF) plot for consensus clustering visualizing the distribution of consensus values across different K values. c) Relative change in the area under the curve (RCSI) quantifies how much clustering stability improves as K increases. K = Number of clusters.*

Supplementary Figure 4. APOE ε4 associations with brain change as a function of age.

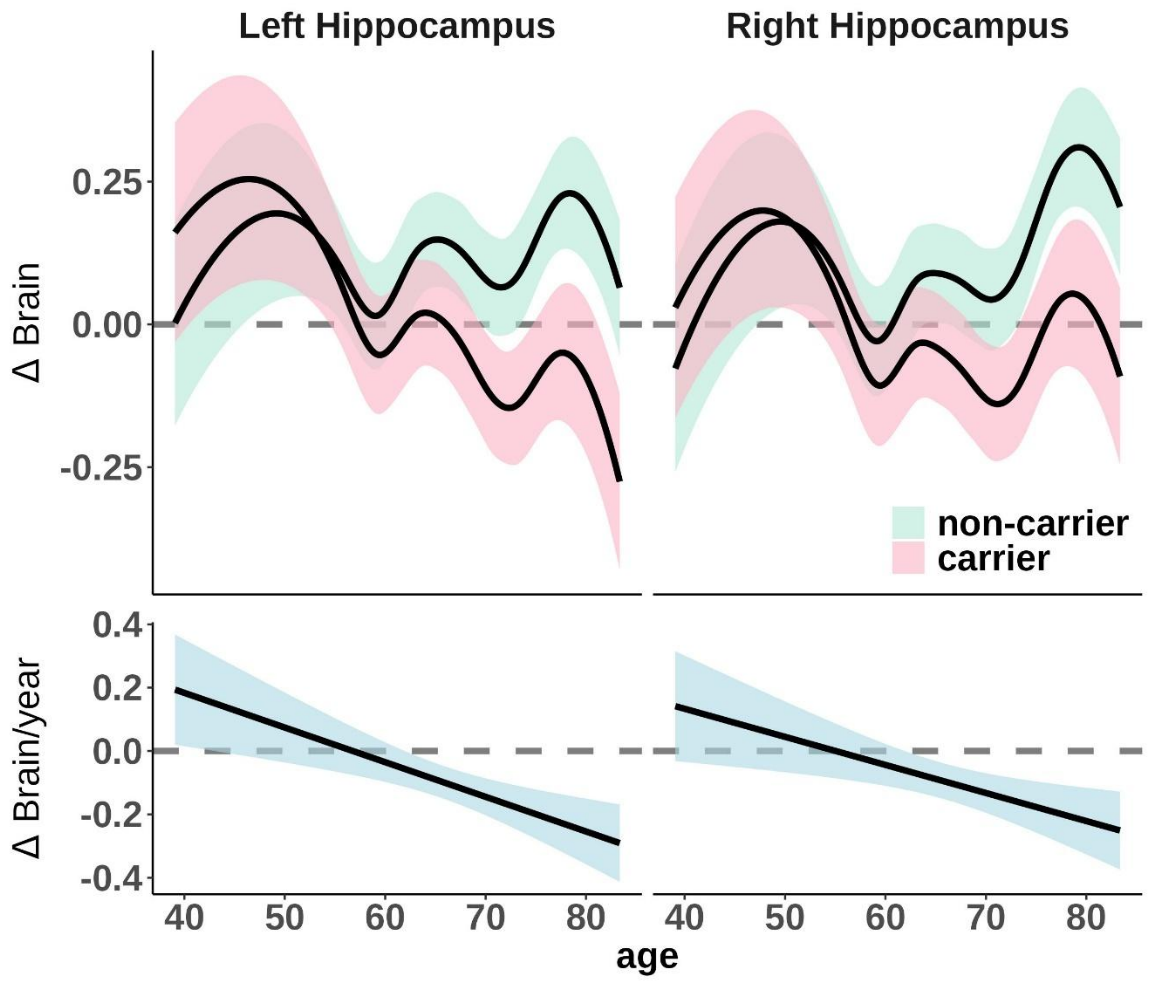


***Supplementary Figure 4. APOE ε4 associations with brain change as a function of age.*** *Upper panel: association between APOE ε4 (carriers, non-carriers) and brain change as a function of age in the left and right hippocampus ($p_{unc} < 0.05$). Note that age and, sex trends are removed based on normative data; thus, brain change should be approximately 0 across age. Lower panel: estimate and 95% CIs for the smooth term of APOE ε4 (i.e., additive effect to the reference smooth trajectories of age in APOE ε4 non-carriers.*

#### Supplementary Table 1. Significant association between brain change and memory change.

| ***Region*** | $\boldsymbol{\beta_w}$ | ***-log10(p)*** | ***-log10(p_fdr)*** |
|---|---|---|---|
| *Left-Amygdala* | 0.155 | 3.699 | 3.699 |
| *Left-Hippocampus* | 0.165 | 3.699 | 3.699 |
| *Left-Inf-Lat-Vent* | 0.132 | 3.699 | 3.699 |
| *lh_S_pericallosal* | 0.127 | 3.699 | 3.699 |
| *Right-Hippocampus* | 0.168 | 3.699 | 3.699 |
| *lh_G_oc-temp_med-Parahip* | 0.131 | 3.097 | 1.722 |
| *Right-Amygdala* | 0.132 | 3.097 | 1.722 |
| *rh_G_Ins_lg_and_S_cent_ins* | 0.135 | 3.000 | 1.683 |
| *Right-Putamen* | 0.136 | 2.796 | 1.541 |
| *Left-Caudate* | 0.074 | 2.745 | 1.541 |
| *rh_G_rectus* | 0.093 | 2.699 | 1.541 |
| *rh_G_temporal_inf* | 0.097 | 2.620 | 1.541 |
| *lh_G_temporal_inf* | 0.091 | 2.585 | 1.541 |
| *rh_G_subcallosal* | 0.114 | 2.585 | 1.541 |
| *rh_S_collat_transv_ant* | 0.112 | 2.585 | 1.541 |
| *Left-Thalamus-Proper* | 0.135 | 2.468 | 1.478 |
| *rh_S_temporal_transverse* | 0.081 | 2.468 | 1.478 |
| *rh_G_temp_sup-Plan_polar* | 0.049 | 2.268 | 1.326 |
| *rh_G_pariet_inf-Supramar* | 0.095 | 2.267 | 1.326 |

*$\beta_w$ = Weighted betas. They were estimated as the density-weighted mean correlation between Δbrain and Δmemory in brain decliners, using the derivative function. $p_{FDR}$ = FDR-corrected p-values. P-values were obtained using wild bootstrapping as out-of-the-box GAMM models are anticonservative[53]. See also **Figure 1** and [Supporting app](#) for complete stats.*

#### Supplementary Table 2. Significant moderator effect of age on association between brain change and memory change.

| ***Region*** | ***-log10(p)*** | ***-log10($p_{fdr}$)*** |
|---:|---:|---:|
| *Right-Inf-Lat-Vent* | 3.699 | 3.699 |
| *Left-Hippocampus* | 3.389 | 1.771 |
| *lh_G_insular_short* | 3.397 | 1.771 |
| *Right-Hippocampus* | 3.391 | 1.771 |
| *Right-Caudate* | 3.096 | 1.575 |
| *Right-Putamen* | 2.990 | 1.548 |
| *Left-Lateral-Ventricle* | 2.691 | 1.316 |

*$p_{FDR}$ = FDR-corrected p-values. P-values were obtained using wild bootstrapping as out-of-the-box GAMM models are anticonservative[53]. See also **Figure 2** and **Supporting app**.*

**Supplementary Table 3**. Cluster assignment and regional PCA loadings.

| Id | Region | PCA1 | PCA2 | PCA33 | PCA4 | Cluster |
|---|---|---|---|---|---|---|
| **1** | Left-Hippocampus | 0.334 | 0.101 | -0.434 | 0.026 | 1 |
| **2** | Right-Hippocampus | 0.324 | 0.111 | -0.406 | 0.033 | 1 |
| **3** | rh_G_temporal_inf | 0.284 | -0.374 | 0.067 | 0.274 | 4 |
| **4** | lh_G_temporal_inf | 0.272 | -0.306 | 0.125 | 0.152 | 4 |
| **5** | Left-Inf-Lat-Vent | 0.260 | -0.066 | -0.487 | -0.125 | 1 |
| **6** | rh_G_temp_sup-Plan_polar | 0.255 | -0.050 | 0.197 | -0.249 | 6 |
| **7** | lh_G_oc-temp_med-Parahip | 0.249 | -0.133 | 0.210 | -0.165 | 8 |
| **8** | Left-Thalamus-Proper | 0.248 | 0.304 | -0.044 | 0.220 | 2 |
| **9** | Right-Amygdala | 0.232 | 0.214 | 0.061 | 0.206 | 3 |
| **10** | rh_S_collat_transv_ant | 0.219 | -0.150 | 0.279 | -0.202 | 8 |
| **11** | rh_G_pariet_inf-Supramar | 0.217 | -0.339 | 0.083 | 0.330 | 4 |
| **12** | rh_G_Ins_lg_and_S_cent_ins | 0.205 | -0.014 | 0.008 | -0.246 | 6 |
| **13** | rh_G_rectus | 0.194 | -0.082 | 0.207 | -0.067 | 5 |
| **14** | Left-Amygdala | 0.188 | 0.104 | 0.087 | 0.129 | 3 |
| **15** | Left-Caudate | 0.188 | 0.427 | 0.234 | 0.122 | 2 |
| **16** | Right-Putamen | 0.163 | 0.470 | 0.314 | 0.101 | 2 |
| **17** | rh_G_subcallosal | 0.141 | 0.039 | -0.002 | -0.325 | 5 |
| **18** | rh_S_temporal_transverse | 0.120 | -0.095 | 0.111 | -0.232 | 7 |
| **19** | lh_S_pericallosal | 0.109 | 0.114 | -0.009 | -0.537 | 5 |

*Loadings from the first 4 components and cluster assignment for those with significant ($p_{FDR}$ < .05) Δmemory – Δbrain associations. See also* ***Figure 3b, c****.*

#### **Supplementary Table 4**. Regional contributions to memory decline

| *cluster* | *Basic β(p)* | *Hippocampus-controlled β(p)* | *Global decline-controlled β(p)* | *All clusters in one model β(p)* |
|---|---|---|---|---|
| *#1* | 0.19(<.001) | -- | 0.08(.13) | 0.09(.001) |
| *#2* | 0.11(.001) | 0.03(.35) | -0.01(.79) | 0.01(.60) |
| *#3* | 0.20(<.001) | 0.11(0.01) | 0.09(.05) | 0.08(.03) |
| *#4* | 0.13(.001) | 0.04(.39) | -0.07(.02) | -0.01(.74) |
| *#5* | 0.23(<.001) | 0.16(.002) | 0.11(.01) | 0.11(0.01) |
| *#6* | 0.09(<.001) | 0.06(.01) | 0.03(.49) | 0.04(.10) |
| *#7* | 0.076(<.001) | 0.05(0.02) | 0.03(.01) | 0.03(.04) |
| *#8* | 0.16(<.001) | 0.10(0.03) | 0.04(.20) | 0.07(.02) |

*Regional contributions with memory decline. Δmemory – Δbrain associations for the different clusters (K = 8). The different analyses were conducted with weighted GAMMs with dataset as random effect. "Basic" includes only the Δbrain cluster term of interest. In the "hippocampus and global-decline-controlled" models, an extra regressor was added, either the hippocampal cluster (#1) or the global decline as estimated from the PCA. In "all clusters in one" model, we included the eight Δbrain clusters in a single GAMM. All p-values except for the last analysis were obtained using bootstrapping. The p-values from the last analysis were obtained as implemented in mgcv R-package*[48], as bootstrapping was extremely conservative, possibly due to cross-correlation between clusters. *See also **Figure 3b, c**. $\beta_w$ = Weighted betas.*

#### Supplementary Table 5. Initial MRI sample

| Dataset | Subjects | Age | | Obs. Memory | | | Time | |
|---|---|---|---|---|---|---|---|---|
| | N (m) | M (SD) | range | N | M (SD) | Range | M (SD) | Range |
| adni | 356 (149) | 74.0 (6.9) | 57.0 -95.2 | 1206 | 3.4 (2.2) | 1 - 12 | 2.6 (2.4) | 0 – 9.6 |
| aibl | 485 (209) | 73.1 (6.1) | 60.0 -92.0 | 871 | 1.8 (1.3) | 1 - 5 | 1.3 (2.1) | 0 – 8.0 |
| bbhi | 930 (477) | 53.5 (7.1) | 40.8 – 67.3 | 1201 | 1.3 (0.5) | 1 - 2 | 0.7 (1.1) | 0 – 3.0 |
| habs | 281 (115) | 75.5 (6.1) | 64.7 – 90.8 | 673 | 2.4 (0.9) | 1 - 4 | 3.5 (2.1) | 0 – 8.5 |
| mpib | 430 (266) | 62.0 (17.2) | 24.8 – 82.0 | 753 | 1.8 (0.5) | 1 - 3 | 1.3 (1.0) | 0 – 3.1 |
| oasis3 | 974 (417) | 70.5 (8.7) | 42.7 – 97.1 | 2120 | 2.2 (1.5) | 1 - 8 | 3.0 (3.9) | 0 – 15.8 |
| ous | 113 (53) | 76.3 (6.2) | 65.7 – 92.7 | 423 | 3.7 (2.0) | 1 - 8 | 5.0 (3.2) | 0- 9.5 |
| preventad | 305 (91) | 64.1 (5.0) | 55.2 – 85.0 | 1360 | 4.5 (1.8) | 1 - 7 | 2.2 (1.5) | 0 – 4.7 |
| ub | 288 (103) | 68.9 (7.0) | 38.3 - 89.6 | 421 | 1.5 (0.8) | 1 - 3 | 1.0 (1.7) | 0 - 5.0 |
| uio | 1179 (402) | 43.6 (19.7) | 20.0 – 93.4 | 2148 | 1.8 (1.2) | 1 - 7 | 2.0 (3.1) | 0 – 11.5 |
| ukb | 1287 (627) | 63.8 (7.1) | 48.2 – 80.6 | 2574 | 2.0 (0) | 2 - 2 | 2.3 (0.1) | 2 – 2.7 |
| umu | 303 (150) | 62.7 (13.6) | 25.0 – 85.0 | 587 | 1.9 (0.8) | 1 - 3 | 3.9 (3.3) | 0 – 9.0 |
| vetsa | 731 (731) | 61.7 (4.5) | 51.1- 71.7 | 1453 | 2.0 (0.8) | 1 - 3 | 6.1 (4.9) | 0 – 13.4 |
| **all** | **7662 (3790)** | **61.8 (14.7)** | **20.0 – 97.1** | **15790** | **2.1 (1.3)** | **1 - 12** | **2.5 (3.1)** | **0 – 15.8** |

*Initial MRI sample sociodemographic and observational details. This sample includes both cross-sectional and longitudinal data and was used for MRI data harmonization, i.e., normative modelling N = Total number of individuals or observations. m = number of males in the sample. NC = Non-carriers. C = Carriers. M = mean. SD = Standard Deviation. Obs. = Observations*

#### Supplementary Table 6. Initial memory sample

| Dataset | Subjects | Age | | Obs. Memory | | | Time | |
|---|---|---|---|---|---|---|---|---|
| | N (m) | M (SD) | Range | N | M (SD) | Range | M (SD) | Range |
| adni | 904 (405) | 74.0 (6.7) | 55.1 – 92.3 | 3824 | 4.2 (2.9) | 1 - 15 | 3.4 (3.2) | 0 – 15.0 |
| aibl | 615 (270) | 73.6 (6.2) | 60.0 – 94.0 | 1239 | 2.0 (1.3) | 1 - 5 | 1.6 (2.2) | 0 – 7.0 |
| bbhi | 973 (500) | 53.5 (7.2) | 29.5 – 67.3 | 1276 | 1.3 (0.5) | 1 - 2 | 0.7 (1.1) | 0 – 3.0 |
| habs | 287 (118) | 75.5 (6.1) | 64.5 – 90.7 | 1251 | 4.4 (2.0) | 1 - 6 | 3.5 (2.1) | 0 – 8.5 |
| mpib | 1899 (936) | 64.0 (16.9) | 23.6 – 91.2 | 3133 | 1.6 (0.7) | 1 - 3 | 2.7 (2.6) | 0 – 6.5 |
| oasis3 | 1067 (463) | 71.4 (8.6) | 45.7 – 96.2 | 6561 | 6.1 (4.6) | 1 - 30 | 7.0 (5.5) | 0 – 31.6 |
| ous | 114 (54) | 76.1 (6.2) | 67.0 – 92.6 | 667 | 5.9 (1.8) | 1 - 7 | 5.2 (1.9) | 0- 6.9 |
| preventad | 306 (91) | 64.4 (5.1) | 55.2 – 85.9 | 1057 | 3.5 (1.8) | 1 - 6 | 2.1 (1.5) | 0 – 4.5 |
| ub | 161 (57) | 69.3 (5.2) | 51.3 – 85.8 | 298 | 1.9 (0.9) | 1 - 3 | 1.8 (2.0) | 0 - 5.2 |
| uio | 939 (303) | 39.7 (17.5) | 20.0 – 93.4 | 1442 | 1.5 (0.8) | 1 - 4 | 2.1 (3.2) | 0 – 11.5 |
| ukb | 33890 (16456) | 65.3 (7.6) | 47.3 – 82.8 | 36520 | 1.1 (0.3) | 1 - 2 | 0.2 (0.6) | 0 – 3.0 |
| umu | 337 (168) | 57.1 (11.4) | 25.0 – 87.9 | 1563 | 4.6 (1.9) | 1 - 7 | 17.6 (8.9) | 0 – 29.0 |
| vetsa | 1592 (1592) | 61.5 (3.9) | 52.1 – 71.1 | 3617 | 2.3 (0.8) | 1 - 3 | 7.6 (4.8) | 0 – 14.4 |
| **all** | **43084 (21413)** | **64.8 (9.7)** | **20.0 – 96.2** | **62448** | **1.4 (1.4)** | **1 - 30** | **1.1 (3.0)** | **0 – 31.6** |

*Initial memory sample sociodemographic and observational details. This sample includes both cross-sectional and longitudinal data and was used for episodic memory data harmonization, i.e., normative modelling N = Total number of individuals or observations. m = number of males in the sample. NC = Non-carriers. C = Carriers. M = mean. SD = Standard Deviation. Obs. = Observations*

#### Supplementary Table 7. Data availability

| *Sample* | *Link* | *PI and/or Admin Contact* | *IRB* |
|---|---|---|---|
| | | *Longitudinal Aging Dataset* | |
| *ADNI* | https://adni.loni.usc.edu/ (O) | Weiner MW; (PI) (AC) | Approved by the Institutional Review Boards of all of the participating institutions |
| *AIBL* | https://aibl.csiro.au/research/ (O) | Christopher Rowe; (PI) | Institutional ethics committees of Austin Health, StVincent's Health, Hollywood |
| *BASE-II* | https://www.base2.mpg.de/en (R) | Lindenberger U, Düzel E, Kühn S (PI); Ludmila Muller | Ethics Committee of the Max-Planck-Institute |
| *BBHI* | https://bbhi.cat/en (R) | Alvaro Pascual-Leone(PI); (AC) | Research institutional review board of the Institut Guttmann |
| *BETULA* | http://www.ufbi.umu.se/english (R) | Lars Nyberg; (PI) | Regional Ethical Vetting Board at Umeå University |
| *COGNORM* | https://www.med.uio.no/klinmed/english/research/groups | Leiv Otto Watne (PI); Anders Martin Fjell; (PI) | Norwegian Regional Committees for Medical and Health Research Ethics and the Data Protector Officer at Oslo University Hospital |

| | | | |
|---|---|---|---|
| | /delirium/index.html; http://www.oslobrains.no (R) | | |
| *HABS* | https://habs.mgh.harvard.edu (O) | Reisa Sperling; (PI); (AC) | Partners Healthcare Human Research Committee |
| *LCBC* | http://www.oslobrains.no (R) | Kristine B. Walhovd; (PI) | Norwegian Regional Committee for Medical and Health Research Ethic; Regional Ethical Committee of South Norway |
| *OASIS3* | https://www.oasis-brains.org/ (O) | Pamela J. LaMontagne; (PI); Daniel Marcus; (PI); https://www.oasis-brains.org/#contact (AC) | Institutional Review Board of Washington University School of Medicine |
| *preventAD* | https://prevent-alzheimer.net; https://openpreventad.loris.ca/ (O) | Jennifer Tremblay; (PI); https://openpreventad.loris.ca/contact/ (AC) | The McGill Institutional Review Board and the Douglas Mental Health University Institute Research Ethics Board |
| *UB* | http://www.ub.edu/bbslab/bbslab/ (R) | David Bartrés-Faz; (PI) | Comisión de Bioética de la Universidad de Barcelona and Hospital Clinic |
| *UKB* | https://www.ukbiobank.ac.uk/ (O) | Rory Collins (PI); (AC) | Northwest Multi-Center Research Ethics Committee [MREC]; https://www.ukbiobank.ac.uk/the-ethics-and-governance-council |
| *VETSA* | https://www.vetsatwins.org/ (O | William S. Kremen(PI); | VETSA and VET Registry Data Security Policies, UCSD Human Subject Committee |

| | | |
|---|---|---|
| | | https://www.vetsatwins.org/for-researchers/ (AC) |

*Data availability, contact and principal investigator information, and ethical approval for the different datasets used. PI = Principal Investigator. AC = Administrative contact. IRB = Institutional Review Boards. O = Openly available. Automatic or semi-automatic data agreements. Fees may apply (e.g. UKB). R = Restricted. Ad-hoc permission is required. Contact PI or AC for specific details on securing access to data.*

#### Supplementary Table 8. Memory measures.

| Dataset | *Subject (Obs.)* | Memory Tests |
|---|---|---|
| **ADNI** | *904 (3824)* | *ADNI-MEM*[1] |
| **AIBL** | *615 (1239)* | *Logical memory short delay recall* |
| | | *Logical memory long delay recall* |
| **BASE-II** | *1899 (3133)* | *VLMT short delay recall* |
| | | *VLMT long delay recall* |
| | | *VLMT learning (sum across trials)* |
| | | *Scene encoding task (hit – fa)*[2] |
| | | *Face-profession task (hits – new fa)*[2] |
| | | *Face-profession task (hits – rear fa)*[2] |
| | | *Object location task (short delay recall)*[2] |
| **BBHI** | *966 (1266)* | *RAVLT learning (sum across trials)* |
| | | *RAVLT long delay recall* |
| | | *RAVLT short delay recall* |
| **BETULA** | *337 (1563)* | *Recall of sentences*[3] |
| **COGNORM** | *114 (667)* | *CERAD short delay recall* |
| | | *CERAD long delay recall* |
| **HABS** | *615 (1239)* | *Logical memory short delay recall* |
| | | *Logical memory long delay recall* |
| | | *SRT delayed recall* |
| | | *SRT total recall* |
| | | *FCsrt free recall* |
| **LCBC** | *938 (1442)* | *CVLT short delay recall* |
| | | *CVLT long delay recall* |
| | | *CVLT learning (sum across trials)* |
| **OASIS3** | *1067 (6561)* | *Logical memory immediate* |
| | | *Logical memory delayed* |
| | | *SRT delayed recall* |
| | | *SRT total recall* |
| | | *WMS associate learning summary score* |
| **preventAD** | *306 (1057)* | *RBANS list recall* |
| | | *RBANS list learning (sum across trials)* |

| | | |
|---|---|---|
| | | *RBANS story immediate memory* |
| | | *RBANS story delayed recall* |
| | | *RBANS figure recall total score* |
| **UB** | *161 (298)* | *RAVLT learning (sum across trials)* |
| | | *RAVLT long delay recall* |
| **UKB** | *33,890 (36,520)* | *PAL[4]* |
| **VETSA** | *1592 (3617)* | *CVLT short delay recall* |
| | | *CVLT long delay recall* |
| | | *CVLT learning (sum across trials)* |

*Memory measures selected in each dataset. A PC was estimated on the first time point when multiple memory measures were available. Subs (Obs.). Subjects and Observations with memory from the initial MRI sample. MMSE = Mini-mental State Examination. RAVLT = Rey Auditory Verbal Learning Test; CVLT = California Verbal Learning Test; PAL = Paired associate learning (#20197 UKB field); Logical memory = Memory subtest of the Wechsler Memory Scale. CERAD = Consortium to Establish a Registry for Alzheimer's Disease (CERAD) Word List Memory test. ADAS = Alzheimer Disease Assessment Scale. VMLT = Verbal Learning and Memory test. SRT = Buschke Selective Reminding Task. RBANS = Repeatable Battery for Assessment of Neuropsychological Status. Story recall = Story recall and recognition task of episodic memory from Wechsler Neuropsychological Battery. FRsrt = Free and Cued selective reminding test. WMS = Weschler Memory scale. FA = False alarms. [1]ADNI-MEM score was computed developed by* [54] and consists of a composite score of memory which includes measures from RAVLT (learning trials, list, recognition and recalls), ADAS (learning trials, recall, and recognitions), *MMSE words, and Logical memory. [2]See* [5,6] *for more details. [3]See* [21] *for more details. [4]See* [*https://biobank.ndph.ox.ac.uk/showcase/refer.cgi?id=2561*](https://biobank.ndph.ox.ac.uk/showcase/refer.cgi?id=2561) *for more information on PAL.*

#### Supplementary Table 9. Scanner acquisition parameters.

| *Dataset* | *Scanner* | *Field* | *Sequence parameters* |
|---|---|---|---|
| *ADNI* | Multisite (n > 50) | 1.5/ 3.0 | See https://adni.loni.usc.edu/methods/documents/mri-protocols/ |
| *AIBL* | Avanto Siemens | 1.5 | MPRAGE. TR: 2300 ms; TE: 2.98 ms, TI: 900 ms; flip angle 9°, slice thickness: 1.25 mm, FoV: 240 x 256, 160 slices. |
| | Verio Siemens | 3.0 | MPRAGE. TR: 2300 ms; TE: 2.98 ms, TI: 900 ms; flip angle 9°, slice thickness: 1.25 mm, FoV: 240 x 256, 160 slices. |
| | Tim Trio Siemens | 3.0 | MPRAGE. TR: 2300 ms; TE: 2.98 ms, TI: 900 ms; flip angle 9°, slice thickness: 1.25 mm, FoV 240 x 256, 160 slices. |
| *BASE-II* | Tim Trio Siemens | 3.0 | MPRAGE. TR: 2500 ms; TE: 4.77 ms, TI: 1100 ms; flip angle 7°, slice thickness: 1.0 mm, FoV 256 x 256, 176 slices. |
| *BBHI* | MAGNETOM Prisma Siemens | 3.0 | MPRAGE. TR: 2400 ms; TE: 2.22 ms, TI: 1000 ms; flip angle 8°, slice thickness: 1.0 mm, FoV 250 x 250, 208 slices. |
| *BETULA* | Discovery GE | 3.0 | 3D FSPGR. TR: 8.19 ms; TE: 3.2ms, TI: 450 ms; flip angle 12°, slice thickness: 1 mm, FoV: 250 x 250, 180 slices. |
| *COGNORM* | Siemens Avanto | | MPRAGE. TR: 2400 ms; TE: 3.79 ms, TI: 1000 ms; flip angle 8°, slice thickness: 1.2 mm, FoV 240 x 240, 160 slices. |
| | Siemens Prisma | 3.0 | MPRAGE. TR: 2400 ms; TE: 2.22 ms, TI: 1000 ms; flip angle 8°, slice thickness: 0.8 mm, FoV 240 x 256, 208 slices, iPat = 2. |
| *HABS*[c] | Tim Trio Siemens | 3.0 | MPRAGE. TR: 2300 ms; TE: 2.98 ms, TI: 900 ms; flip angle 9°, slice thickness: 1.2 mm, FoV 240 x 256, 160 slices. MPRAGE. TR: 2200 ms; TE: 1.5/3.4/5.2/7.0 ms, TI: 1100 ms; flip angle 7°, slice thickness: 1.2 mm, FoV: 228 x 228, 144 slices, Multi-echo = x4. |
| *LCBC* | Avanto Siemens | 1.5 | MPRAGE. TR: 2400 ms; TE: 3.79 ms, TI: 1000 ms; flip angle 8°, slice thickness: 1.2 mm, FoV 240 x 240, 160 slices. |
| | Siemens Skyra | 3.0 | MPRAGE. TR: 2300 ms; TE: 2.98 ms, TI: 850 ms; flip angle 8°, slice thickness: 1 mm, FoV: 256 x 256, 176 slices. |
| | Siemens Prisma | 3.0 | MPRAGE. TR: 2400 ms; TE: 2.22 ms, TI: 1000 ms; flip angle 8°, slice thickness: 0.8 mm, FoV 240 x 256, 208 slices, iPat = 2. |

| | | | |
|---|---|---|---|
| *OASIS3* | Siemens Vision | 1.5 | MPRAGE. TR: 9,7 ms; TE: 4.0 ms, TI: 20 ms; flip angle 10°, slice thickness: 1.25 mm, FoV: 256 x 256, 160 slices. |
| | Siemens Sonata | 1.5 | MPRAGE. TR: 9,7 ms; TE: 3.9 ms, TI: 20 ms; flip angle 15°, slice thickness: 1 mm, FoV: 224 x 256, 160 slices. |
| | Tim Trio Siemens | 3.0 | MPRAGE. TR: 2400 ms; TE: 3.1 ms, TI: 1000 ms; flip angle 8°, slice thickness: 1 mm, FoV 256 x 256, 176 slices. |
| | Magnetom Vida Siemens | 3.0 | MPRAGE. TR: 2300 ms; TE: 2.3 ms, TI: 900 ms; flip angle 9°, slice thickness: 1.2 mm, FoV 240 x 256, 176 slices. |
| | Siemens BioGraph mMR | 3.0 | MPRAGE. TR: 2300 ms; TE: 2.3 ms, TI: 900 ms; flip angle 9°, slice thickness: 1.2 mm, FoV 240 x 256, 176 slices. |
| *preventAD* | Tim Trio Siemens | 3.0 | MPRAGE. TR: 2300 ms; TE: 2.98 ms, TI: 900 ms; flip angle 9°, slice thickness: 1 mm, FoV 240 x 256, 176 slices. |
| *UB* | Tim trio Siemens | 3.0 | MPRAGE. TR: 2400 ms; TE: 2.98 ms, TI: 900 ms; flip angle 9°, slice thickness: 1 mm, FoV: 256 x 256, 240 slices. |
| *UKB* | Siemens Skyra[a] | 3.0 | MPRAGE. TR: 2000 ms; TE: - ms, TI: 880 ms; flip angle -, slice thickness: 1 mm, FoV: 208 x 256, 256 slices. |
| | Siemens Avanto | 1.5 | MPRAGE. TR: 1000 ms; TE: 3.31 ms, TI: 1000 ms; flip angle 7°, slice thickness: 1.33 mm, FoV 256 x 256, 128 slices |
| | Siemens Symphony | 1.5 | MPRAGE. TR: 1000 ms; TE: 3.31 ms, TI: 1000 ms; flip angle 7°, slice thickness: 1.33 mm, FoV 256 x 256, 128 slices |
| *VETSA* | Discovery 750× GE[d] | 3.0 | 3D FSPGR. TR: 8.084 ms; TE: 3.164ms, TI: 600 ms; flip angle 8°, slice thickness: 1.2 mm, FoV: 256 x 256, 176 slices. |
| | Tim Trio Siemens | 3.0 | MPRAGE. TR: 2170 ms; TE: 4.33 ms, TI: 1100 ms; flip angle 7°, slice thickness: 1.2 mm, FoV 256 x 256, 160 slices. |

*Scanner Parameters for each dataset. TR = Repetition Time; TE = Echo Time; TI = inversion time; FoV = Field of View, iPat = in-plane acceleration. [a,c,d]Two matched scanners. [b]Several matched scanners.*
